# Females drive postmating reproductive trait evolution across *Drosophila* species, but not via remating rate

**DOI:** 10.1101/2024.10.15.618555

**Authors:** Brooke Peckenpaugh, Leonie C. Moyle

## Abstract

While traits that contribute to premating sexual interactions are known to be wildly diverse, much less is known about the diversity of postmating (especially female) reproductive traits and the mechanisms shaping this diversity. To assess the rate, pattern, and potential drivers of postmating reproductive trait evolution, we analyzed male and female traits across up to 30 *Drosophila* species within a phylogenetic comparative framework. In addition to postmating reproductive morphology (e.g., sperm length, reproductive tract length and mass), we also quantified mating behaviors including female remating rate—a common proxy for the strength of postmating sexual selection. We found evidence for strong coevolution between male and female postmating traits (specifically sperm length and sperm storage organ size). However, remating rate was not associated with the rate of evolution or exaggeration of either male or female postmating reproductive morphology, once phylogenetic relatedness was accounted for. We infer that female-mediated and intersexual selection predominantly drive the evolution of our postmating morphological traits, including via divergent male and female interests in controlling paternity. In comparison, remating rate has a complex and likely secondary role in shaping this evolution, in part because this trait can be both a driver and a product of postmating selection.

## 1 Introduction

In sexual organisms, reproductive traits are often extravagantly diverse and rapidly evolving. This diversity extends to traits and genes that act during and after mating, whose rapid evolution has been documented across organisms that range from tomatoes to primates (Moyle et al. 2021, Dorus et al. 2004). These postmating traits have been studied extensively in males. For example, *Drosophila* male genitalia (Hosken and Stockley 2004), seminal fluid protein genes (Wagstaff and Begun 2005, Haerty 2007), and sperm size, form, and function (Immler et al. 2011, Lüpold et al. 2016) have all diverged significantly between even closely related species. For instance, the sperm gigantism of some *Drosophila*—including the 5.83 cm long sperm of *Drosophila bifurca—*is one of the most extreme examples of a sexual ornament in nature (Lüpold et al. 2016). These instances of rapid postmating trait divergence are often proposed to be a consequence of intrasexual selection (i.e., sperm competition) because variation in traits like sperm length or number can affect the number of offspring sired during competitive matings (Parker 1970). Nonetheless, postmating trait evolution could also be driven by intersexual selection, in the form of female postmating preferences (Eberhard 1996), or by sexually antagonistic coevolution between conflicting male and female reproductive strategies (Chapman et al. 2003). However, assessing the relative contributions of these different evolutionary forces to reproductive trait evolution is challenging. To do so requires not only information about male reproductive evolution, but also a matched assessment of female traits, including in contexts where these different types of sexual selection could systematically vary in strength.

Like male traits, female reproductive traits are predicted to play an important role in shaping the outcome of postmating interactions. In *Drosophila*, for example, females can exert control over the paternity of their offspring via differential storage and/or use of sperm from two different kinds of sperm storage organs after mating: the seminal receptacle (SR) and the paired spermathecae (ST) (Manier et al. 2013). The precise mechanisms of this ‘cryptic female choice’ (Eberhard 1996) remain unclear in most species. However, experimental evolution in *Drosophila melanogaster* suggests that SR length itself could be a proximate mechanism for female choice among sperm of different lengths, and that female traits like the SR can exert selection directly on male postmating traits, producing evolutionary change in just dozens of generations (Miller and Pitnick 2002). Evidence for male-female postmating coevolution also comes from significant phylogenetic correlations between male and female trait values, including sperm length and female sperm storage organ length (as observed in the *Drosophila obscura* group; Holman et al. 2008). These observations suggest that female-mediated processes and/or male-female coevolution could also play a substantial role in the evolution of postmating traits among *Drosophila* species, including in the exaggeration of sperm.

Another way to assess the role of sexual selection in driving reproductive trait evolution is by comparing species with different expected intensities of sexual selection. Female remating rate (how rapidly a female remates subsequent to a first mating) is a frequent proxy for the strength of postmating sexual selection because it determines how many males’ sperm overlap in a single female reproductive tract (Markow 2002, Firman et al. 2017). Female remating rate could therefore directly affect the opportunity for both sperm competition and for cryptic female choice, each of which should be more intense with more frequent remating. For example, among 21 *Drosophila* species, frequent female remating (daily or more) is broadly associated with exaggerated ejaculates, suggesting stronger selection for these postmating male traits (Markow 2002); in comparison, infrequent female remating (less than daily) is associated with external male secondary sexual characters (such as sex combs or nuptial gifts) that are instead thought to reflect pre-mating sexual selection (Markow 2002). Similarly, among 13 species in the *Drosophila obscura* group, female remating rate is significantly associated with both male reproductive tract mass and spermathecal area (one of the female sperm storage organs), after accounting for phylogenetic relatedness (Holman et al. 2008). Comparative approaches such as the latter are particularly powerful because they can account for the effect of species relatedness on the pattern and distribution of trait variation, and therefore can identify patterns of trait change and trait associations that are driven by evolutionary processes beyond shared history.

In this study, our goal was to use a quantitative comparative phylogenetic framework to identify the predominant drivers of both male and female postmating trait evolution, including the role of varying strengths of sexual selection between sexes and between different lineages. To do so, we assessed the evolution of male and female reproductive traits across at least 15 and up to 30 *Drosophila* species (whose common ancestor lived roughly 50MYA; Suvorov et al. 2022) that vary in their expected intensity of postmating sexual selection. We made the following predictions based on prior expectations of the strength of selection acting on postmating trait evolution:

1. Reproductive traits, both behavioral and morphological, will evolve more rapidly than non-reproductive traits (i.e., thorax length, body mass).
2. Male postmating traits (e.g., sperm length, testis length, testis color, male reproductive tract [MRT] mass) will generally evolve more rapidly than female traits (e.g., SR length, ST area, ST color, female reproductive tract [FRT] mass), because male traits are subject to both inter- and intrasexual selection while female postmating traits are primarily affected by intersexual selection alone. This comparison therefore provides information on the relative contribution of intrasexual selection to shaping postmating evolution.
3. Male and female postmating trait evolution will be significantly associated, if intersexual selection plays an important role in the evolution of these traits.
4. Faster evolution and/or coevolution in male and female postmating traits will be associated with more rapid female remating, if this trait magnifies the intensity of postmating sexual selection. Similarly, higher remating rates should be associated with more complex and/or energetically costly reproductive morphology (e.g., larger sperm storage organs, larger testes, longer sperm).
5. Female remating rate will be more closely associated with postmating trait evolution than premating or perimating (during mating) trait evolution, if it reflects the intensity of postmating sexual selection.

Our goal was to address five key questions regarding postmating trait evolution among these species: What is the pattern and rate of reproductive trait evolution (prediction 1)? Is there evidence for stronger sexual selection acting on males or females (prediction 2)? Is there evidence for co-evolution of male and female postmating traits (prediction 3)? Is rapid remating associated with the exaggerated (a) size or (b) evolutionary rate of postmating reproductive traits in either males or females (prediction 4)? And, is remating rate more closely associated with postmating trait evolution than premating or perimating trait evolution (prediction 5)? Together, answers to these questions can reveal whether postmating trait evolution is predominantly male- driven, female-driven, or shaped by both sexes, as well as uncover factors that facilitate or constrain the dynamics of their evolution.

## 2 Methods

### 2.1 Data collection for developmental, behavioral, and morphological traits

We collected data for our traits of interest from two sources. First, for individual lines of each of 16 different species (see Table S1), we directly quantified eight measures of reproductive morphology, and for 15 of these lines (all except *D. immigrans*) also quantified five elements of reproductive behavior and development (Table S2, Table S3). All our experimental stocks were reared on standard media (prepared by the Bloomington *Drosophila* Stock Center) and kept at 25°C. Our methodology for collecting these experimental data is described below.

Second, we complemented this dataset by collating previously published data on three reproductive morphology traits and three reproductive behavior and development traits (Table S4). Together, this allowed us to assess patterns of trait evolution across 15 to 30 *Drosophila* species, depending upon the specific trait(s) and relationships analyzed.

### 2.2 Developmental and behavioral data collection

#### 2.2.1 Age of female reproductive maturity

To accurately estimate female mating and remating traits, we first had to experimentally determine the female age of reproductive maturity for each species. We did this using a previously established experimental approach (e.g., Pitnick et al. 1995, Snook and Markow 2001, Holman et al. 2008), which was replicated across ten individuals for each of 15 species. See Supplementary Methods section 2.2.1 for experimental details.

#### 2.2.2 Female remating rate

We quantitatively estimated the female remating rate for each line/species by experimentally determining the minimum interval within which the majority of previously mated females would remate (female intermating interval), based on established methods (e.g., Snook and Markow 2001, Holman et al. 2008). We replicated this across five individuals for each of 15 species. We transformed this variable into the female remating rate by taking the inverse of the intermating interval. See Supplementary Methods section 2.2.2 for experimental details.

### 2.3 Morphological data collection

#### 2.3.1 Thorax length, reproductive tract length, and pigmentation

Using reproductively mature flies (as previously determined experimentally), we measured male and female thorax lengths under a microscope; this is a standard measure of body size in studies of *Drosophila* because it strongly correlates with the size of other characters such as wing length (Robertson and Reeve 1952). We then dissected either seminal receptacles and spermathecae (females) or testes (males) to measure their length and pigmentation. Length and pigmentation measurements were replicated across five individuals for each of 16 species. See Supplementary Methods section 2.3.1 for experimental details.

#### 2.3.2 Body mass and reproductive tract mass

We measured body and reproductive tract mass in reproductively mature flies according to Holman et al. (2008). Mass measurements were replicated across five individuals for each of 16 species. See Supplementary Methods section 2.3.2 for experimental details.

#### 2.3.3 Sperm length

Sperm length was also experimentally assessed in reproductively mature flies for a subset of ten of the species for which other morphological traits were directly measured (specifically, *D. melanogaster, D. simulans, D. yakuba, D. teissieri, D. ananassae, D. pseudoobscura, D. persimilis, D. mojavensis, D. arawakana,* and *D. immigrans*). The five experimental species with longest sperm (*D. virilis, D. lummei, D. americana, D. novamexicana,* and *D. hydei*) were not assessed, as this experimental procedure is not effective for accurately measuring sperm longer than ∼2 mm. We measured sperm length for 5-10 randomly chosen sperm per individual and repeated these measurements across five individuals per species. See Supplementary Methods section 2.3.3 for experimental details.

### 2.4 Previously published trait data

To address our questions in additional species, we collated previously published data on three reproductive morphology traits (SR length, testis length, and sperm length) and three reproductive behavior and development traits (remating rate, mating duration, and female age of reproductive maturity) (Table S4). These additional data were drawn from eight studies (see Table S4; Atkinson 1979, Joly and Bressac 1994, Markow 1996, Pitnick et al. 1995, Pitnick 1996, Pitnick et al. 1999, Markow 2002, Holman et al. 2008). Because the specific methods for measuring female remating rate are not uniform across these studies, in this larger dataset we recoded female remating data into two discrete categories—rapid versus infrequent remating— according to whether more than half of females remated within 24 hours. This specific binary classification has been previously used in analyses of female remating (Markow 1996, Markow 2002).

### 2.5 Phylogeny

We used pruned versions of the *Drosophila* phylogeny from Suvorov et al. (2022), so that they included only the species for which we had matching trait data. We time-calibrated the phylogeny using a discrete clock model, also using node ages from Suvorov et al. (2022).

Specifically, we assumed that extant members of the genus *Drosophila* split from the other Drosophilidae (*Leucophenga, Scaptodrosophila*, and *Chymomyza*) at 53 MYA, the split between the subgenera *Sophophora* and *Drosophila* occurred 47 MYA, and the split within the *Drosophila* subgenus occurred 34 MYA (Suvorov et al. 2022). This, and all subsequent statistical analyses, were carried out in R version 4.3.1.

### 2.6 What is the pattern and rate of reproductive evolution?

#### 2.6.1 Modeling trait evolution

Following Revell and Harmon (2022), we modeled either continuous or discrete character evolution, depending on the specific focal trait. For continuous traits, we fit each log-transformed trait with three models using the fitContinuous function in geiger (Pennell et al. 2014): Brownian motion (BM), early burst (EB), and Ornstein-Uhlenbeck (OU). We compared AIC scores and Akaike weights to identify the best-fitting model. In ambiguous cases, we ran a likelihood-ratio test to determine whether we had sufficient evidence to reject the simpler BM model. After identifying the best-fitting model, we estimated the rate of evolution (σ^2^) and the root state (Z0). We fit remating rate, which is a binary discrete trait, with two M*k* models using the fitDiscrete function in geiger (Pennell et al. 2014): equal-rates (ER) and all-rates-different (ARD). For testis color, which is discrete but not binary, we fit three M*k* models: ER, symmetric transition (SYM), and ARD. We compared AIC scores and Akaike weights to identify the best-fitting model. In ambiguous cases, we ran a likelihood-ratio test to determine whether we had sufficient evidence to reject the simpler ER model. After identifying the best-fitting model for our discrete traits, we estimated the transition rate (*q*) of each.

#### 2.6.2 Ancestral state reconstruction

We estimated ancestral states for our discrete traits as well as any continuous traits that evolve via Brownian Motion. For continuous traits, we estimated ancestral states assuming a BM model of evolution using the fastAnc function in phytools (Revell 2012). We plotted estimated ancestral states on the tree using contMap. For discrete characters, we estimated ancestral states using stochastic character mapping (Huelsenbeck et al. 2003). We generated 1000 stochastic character maps using the make.simmap function, and plotted them using densityMap, both from the phytools package (Revell 2012). For testis color, we estimated ancestral states without specifying a root node state and also reran the analysis with the root node constrained to be orange. This latter model accounts for the possibility that the ability to produce orange testes cannot be regained once it is lost.

### 2.7 Is there evidence for stronger sexual selection acting on males or females?

To maximize our power to address this question, we used our expanded dataset that had quantitative data on both SR and sperm length in 30 species. We compared rates of trait evolution between SR length and sperm length (both log-transformed). To do so, we fit two models of trait evolution using the ratebytree function in phytools (Revell 2012, Adams 2013). In one model, we allowed different evolutionary rates for each trait, and in the other, we constrained them to a common rate of trait evolution. We then conducted a likelihood-ratio test to determine whether the model with multiple rates fit better than the common rate model.

### 2.8 Is there evidence for coevolution between male and female postmating traits?

We assessed this question in both our original dataset (n = 15 species) and our expanded dataset (n = 30 species). To do so, we fit linear regression models to our log-transformed data using phylogenetic generalized least squares (PGLS). We used SR length as the dependent variable, and ran models with both testis length and sperm length as predictors while accounting for phylogenetic relationships. We did this using the gls function in the nlme package (Pinheiro and Bates 2006). To generate the correlation structure of our phylogeny, we used the corPagel function in the ape package (Popescu et al. 2012). The λ parameter is a multiplier of the off- diagonal elements in the expected correlations among species, which allows for either more or less phylogenetic signal than expected under a Brownian motion model (Pagel 1999).

To visualize the effects of controlling for the phylogeny, we also computed phylogenetically independent contrasts (PICs) between testis length and SR length. We did this using the pic function in the ape package.

### 2.9 Is rapid remating associated with the exaggerated (a) size or (b) evolutionary rate of postmating reproductive traits in either males or females?

We assessed these questions using both our original dataset (n = 15 species) and our expanded dataset (n = 29 species). For our original dataset, we fit linear regression models to our log-transformed data using PGLS because our predictor variable (female remating rate) was a continuous trait. We assessed remating rate as a predictor for all measures of postmating reproductive traits (SR length, ST area, ST pigmentation, FRT mass, testis length, testis color, sperm length, and MRT mass); we included a size predictor (thorax length or body mass [with reproductive tract removed] in the relevant sex) as a covariate in each model. To visualize the effects of controlling for the phylogeny, we also computed PICs between both testis and SR length and remating rate.

For our expanded dataset, where remating rate is coded as a discrete character, we instead assessed the correlation between remating rate and reproductive morphology using threshold models (Felsenstein 2005, Felsenstein 2012). Specifically, we assessed the correlation between classes of female remating rate (rapid versus infrequent) and SR length, testis length, and sperm length. We implemented threshold models using the threshBayes function in the phytools package (Revell 2012, Revell 2014).

Finally, using our expanded dataset, we also tested for differences in evolutionary rate or trait correlations in different parts of the phylogeny. Specifically, we tested whether rates of SR or sperm length evolution, or the coevolution of these two traits, differed in lineages of the tree where females remate rapidly compared to lineages where females remate infrequently. To do this, we fit a multivariate Brownian model in which the evolutionary covariance between characters is allowed to be different in different parts of the phylogeny (Revell and Collar 2009). We implemented this analysis using the evolvcv.lite function in the phytools package (Revell 2012, Revell et al. 2022). This allowed us to compare eight possible combinations of models: where rates for either SR length or sperm length, or both, were allowed to vary or not in rapidly remating vs. infrequently remating groups, and for each of these models, whether the coevolution between SR length and sperm length was allowed to be different or not in rapidly remating vs. infrequently remating groups.

### 2.10 Is remating rate more closely associated with postmating trait evolution than premating or perimating trait evolution?

We assessed this question using both our original dataset (n = 15 species) and our expanded dataset (n = 29 species). For our original dataset, we fit linear regression models to our data using PGLS. We assessed remating rate as a predictor for mating latency, mating duration, and age of female reproductive maturity; we included a size predictor (female thorax length) as a covariate in each model. For our expanded dataset, in which remating rate is coded as a discrete character, we assessed the correlation between remating rate and reproductive behavior using threshold models (Felsenstein 2005, Felsenstein 2012). Specifically, we assessed the correlation between female remating rate with mating duration and the age of female reproductive maturity.

## 3 Results

### 3.1 What is the pattern and rate of reproductive evolution?

We found extensive diversity in reproductive morphology across *Drosophila* species, in both females (Figure 1a, Figure S1, Table S2) and males (Figure 1b, Figure S1, Table S3). This included significant variation in gonad size, with seminal receptacle (SR) size ranging nearly two orders of magnitude from 0.480 mm (*D. pseudoobscura*) to 33.014 mm (*D. hydei*) on average, and testes size ranging more than 15-fold from 0.840 mm (*D. persimilis*) to 25.762 mm (*D. hydei*) on average (Table S2 and S3). Spermatheca (ST) area similarly ranged from 0.001 mm^2^ in *D. mojavensis* (which has vestigial, non-functional spermathecae (Pitnick et al. 1999)), to 0.018 mm^2^ on average in *D. virilis*. We also saw variation in pigmentation across reproductive organs. Testes ranged from white to yellow or orange (Figure S1, Table S3), and spermathecae varied in pigment intensity, with *D. mojavensis* being the lightest and *D. persimilis* being the darkest (Figure S1, Table S2).

**Figure 1.**
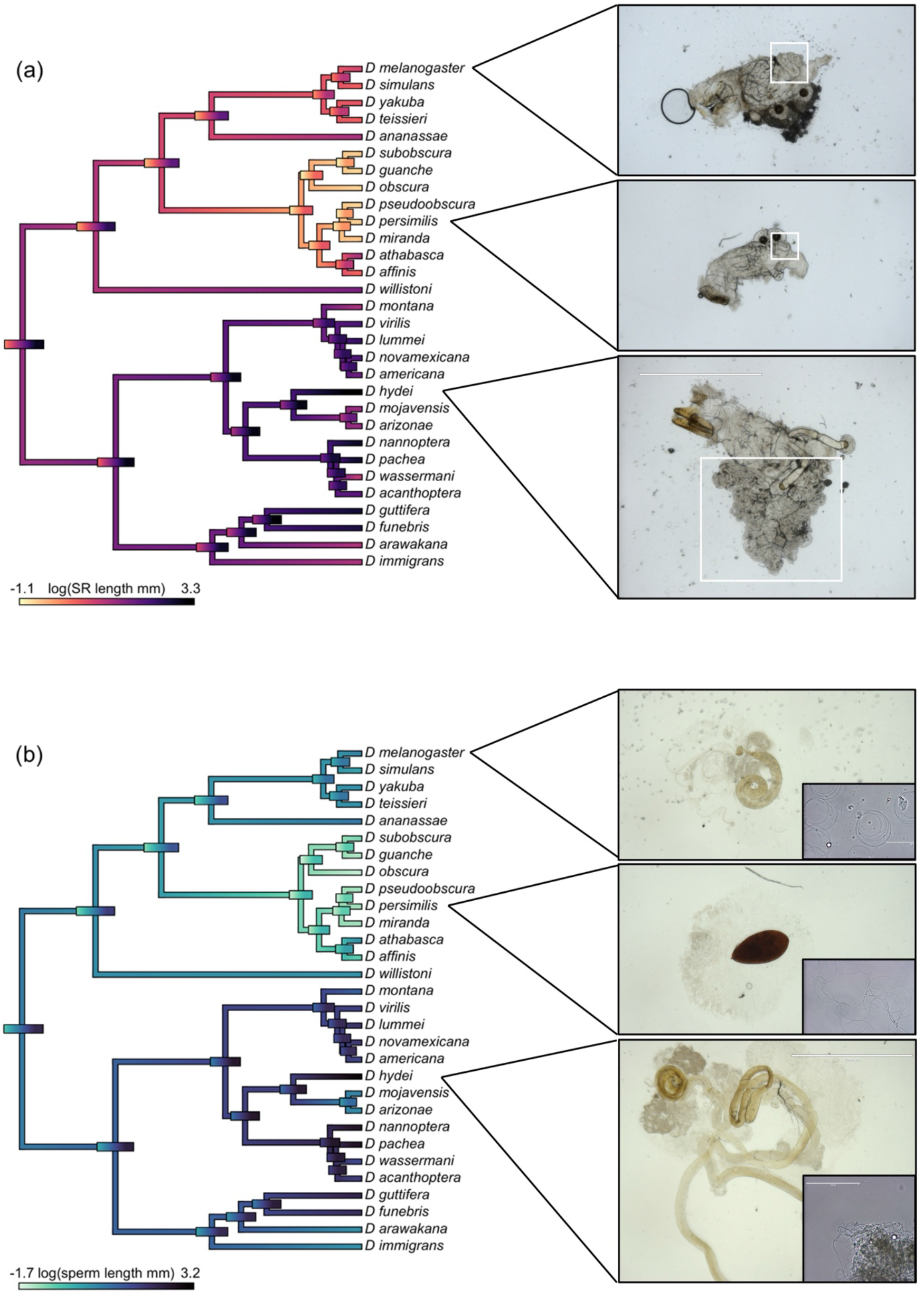
Evolutionary reconstruction of variation in (a) seminal receptacle (SR) length and (b) sperm length (both on a log-scale) across the *Drosophila* phylogeny. The color gradient shows observed (at the tips) or estimated (internal branches) trait values. Error bars show the uncertainty associated with the estimated values for the trait at internal nodes of the tree. For three selected species, photos of the female reproductive tract (a) or testes and sperm (b) are shown. The SR is highlighted with a white box in (a).

Developmental and behavioral reproductive traits also varied substantially between species. For instance, the age of female reproductive maturity ranged from 2 days for *D. melanogaster* and *D. simulans*, to 14 days in *D. virilis* (Table S2). Female remating interval, or the length of time from first mating to second mating, ranged from half a day in *D. hydei* to approximately 23 days in *D. ananassae*. Mating latency (the length of time from setting up a cross until mating begins) was shortest in *D. pseudoobscura*, where it takes only 4 minutes on average, and longest in *D. virilis* where it took about 66 minutes. Mating duration was shortest in *D. hydei*, at only around 2 minutes, and longest in *D. yakuba*, at 38 minutes on average (Table S2).

To evaluate the rates of evolution for each trait, we modeled both continuous (Table S5) and discrete (Table S6) character evolution across the phylogeny. Among all our traits, female and male thorax length had the lowest rates of expected change in variance per million years (σ^2^ = 0.003 and σ^2^ = 0.002, respectively) whereas mating duration had the fastest rate at σ^2^ = 0.245. In general, behavioral traits (including mating latency and duration) were inferred to have among the most rapid rates, and tended to evolve according to a different model of evolution (OU rather than Brownian), compared to the majority of reproductive postmating traits (Table S5); the exceptions among the latter were FRT mass and spermathecal (ST) size, whose rates were the highest among morphological postmating traits and also best fit an OU model of evolution (Table S5). Estimated root states for our traits (Table S5) suggest that the ancestral lineage to all our analyzed species was characterized by relatively infrequent remating (estimated remating interval of 3.73 days), and relatively shorter SR, testis, and sperm length (compared to the mean of our contemporary species; Figures S2, S3, S4). Finally, we estimated ancestral states for the continuous traits that evolve via Brownian motion (Figure 1, Figures S2-S6) and for discrete traits (Figure 4, Figure S7); these provide projections for trait values at internal nodes and branches across the phylogeny.

### 3.2 Is there evidence for stronger sexual selection acting on males or females?

We addressed this question by focusing on SR length (female trait) and sperm length (male trait) because it allowed us to use the largest possible dataset. Across all species, our inferred rates of evolution for SR length and sperm length were σ^2^ = 0.069 and σ^2^ = 0.059, respectively (Table S5). Using our expanded dataset (n = 30 species), we conducted a likelihood-ratio test to determine whether a model with different evolutionary rates for each trait fit our data better than a model with a common rate of trait evolution. We found that the difference was not significant (*p* = 0.642), suggesting that the SR and sperm are not evolving at significantly different rates (Figure 1A and B).

### 3.3 Is there evidence for coevolution between male and female postmating traits?

In our experimental group of 15 *Drosophila* species, we addressed this question using SR length (female trait) and testis length (male trait)—the traits for which we had the most species data. SR length was significantly associated with testis length, both before and after phylogenetic correction (Figure 2, Table S7). First, using ordinary linear models, we found a significant correlation between SR and testis length (OLS, β = 1.295, *t* = 9.841, *p* < 0.001; Figure 2). This association held after we accounted for phylogenetic relationships, both when using phylogenetically independent contrasts (PIC, β = 1.029, *t* = 6.631, *p* < 0.001; Figure 2) and phylogenetic generealized least squares (PGLS, Table S7). We found the same result in our expanded dataset, where SR length was significantly associated with both testis length (n = 20 species) and sperm length (n = 30 species) (Table S8). (This analysis included sperm length as this male trait is the most widely measured in other studies.) These strong associations are consistent with coordinated evolution between male and female traits.

**Figure 2.**
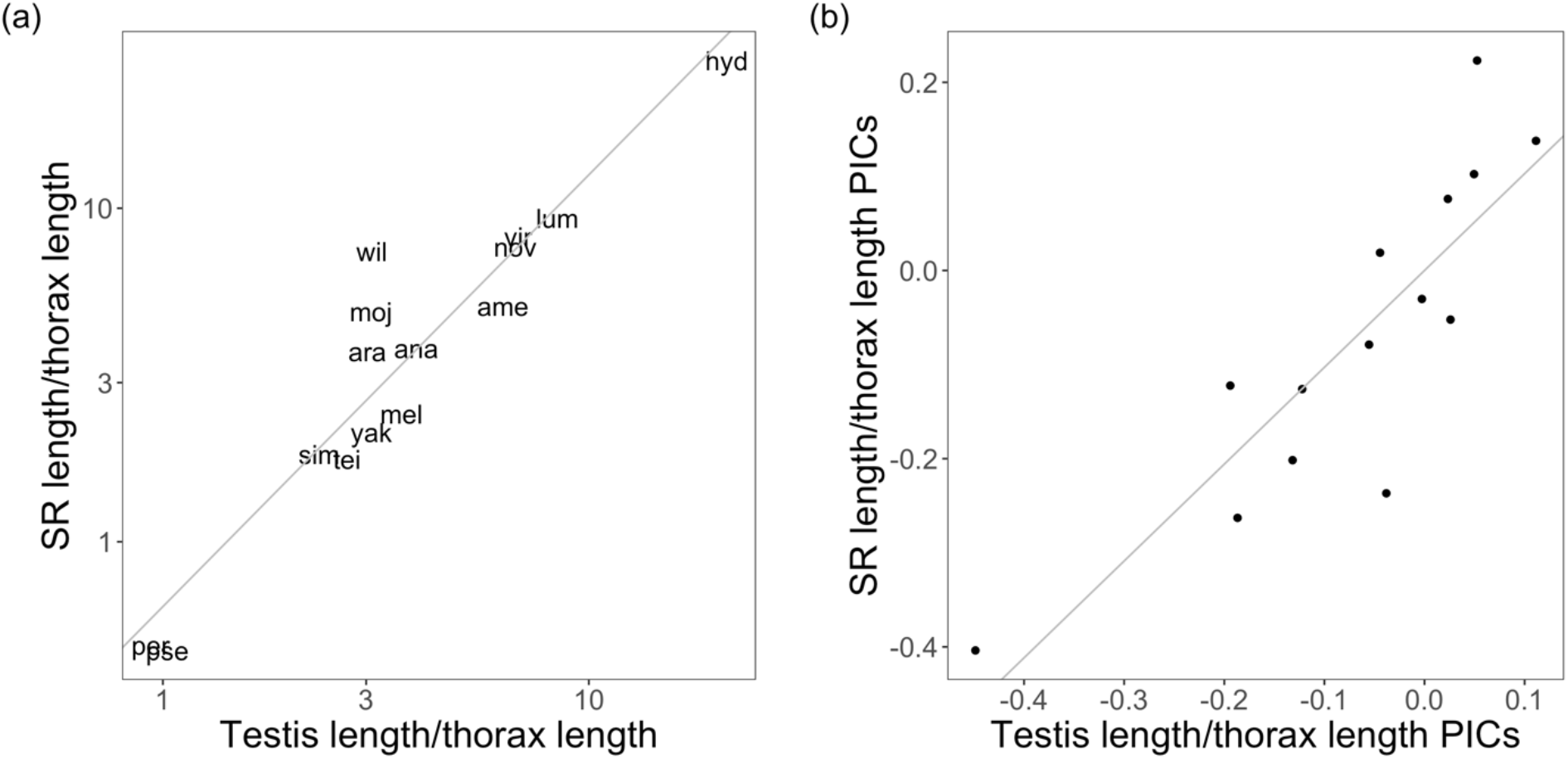
Testis length is correlated with SR length across *Drosophila* species (a), including after performing phylogenetically independent contrasts (PICs) (b). Both axes in (a) are on a log scale. Both testis length and SR length have been corrected for body size by dividing values over thorax length.

### 3.4 Is rapid remating associated with the exaggerated (a) size or (b) evolutionary rate of postmating reproductive traits in either males or females?

First, we looked for a general association between female remating rate and the exaggerated size of postmating reproductive traits. Using ordinary linear models on data that we collected across 15 *Drosophila* species and our quantitative measure of remating rate, we found a significant correlation between female remating rate and testis length (OLS, β = 0.340, *t* = 2.294, *p* = 0.039; Figure 3a), and a marginally significant correlation between female remating rate and SR length (OLS, β = 0.413, *t* = 1.934, *p* = 0.075; Figure 3c). However, when accounting for phylogeny (either using PIC or PGLS), we found no association between female remating rate and either testis or SR length (Figure 3b,d). We similarly found no association between remating rate and most other female and male morphological traits measured, including ST area, ST pigmentation, FRT mass, testis length, testis color, and sperm length (Table S7, Table S9).

**Figure 3.**
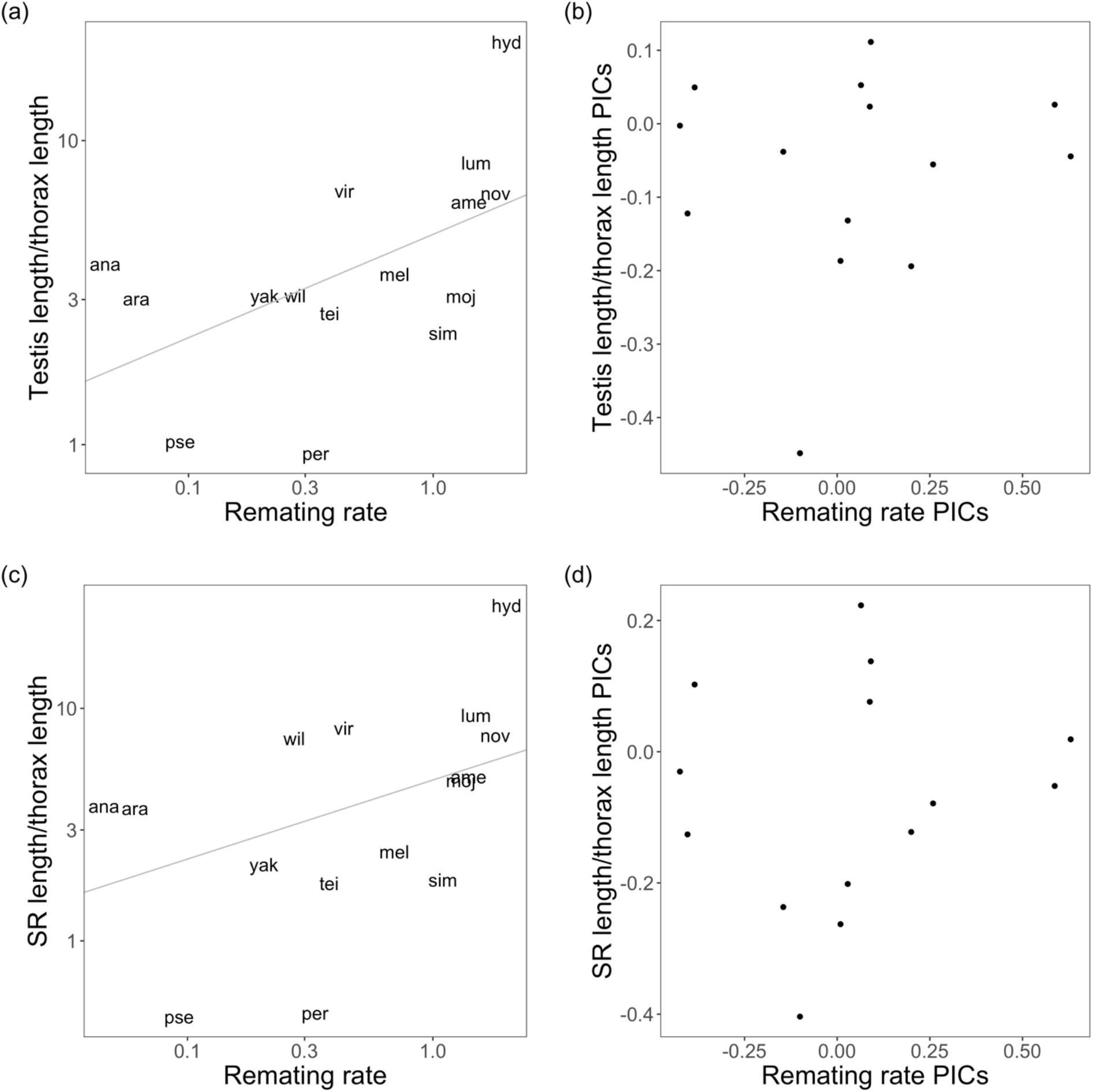
Before accounting for phylogenetic relationships, testis length (a) is associated with remating rate and SR length (c) is marginally associated with remating rate. Using phylogenetically independent contrasts (PICs), neither testis length (b) nor SR length (d) are associated with remating rate. Both axes in (a) and (c) are on a log scale. Both testis length and SR length have been corrected for body size by dividing values over thorax length.

**Figure 4.**
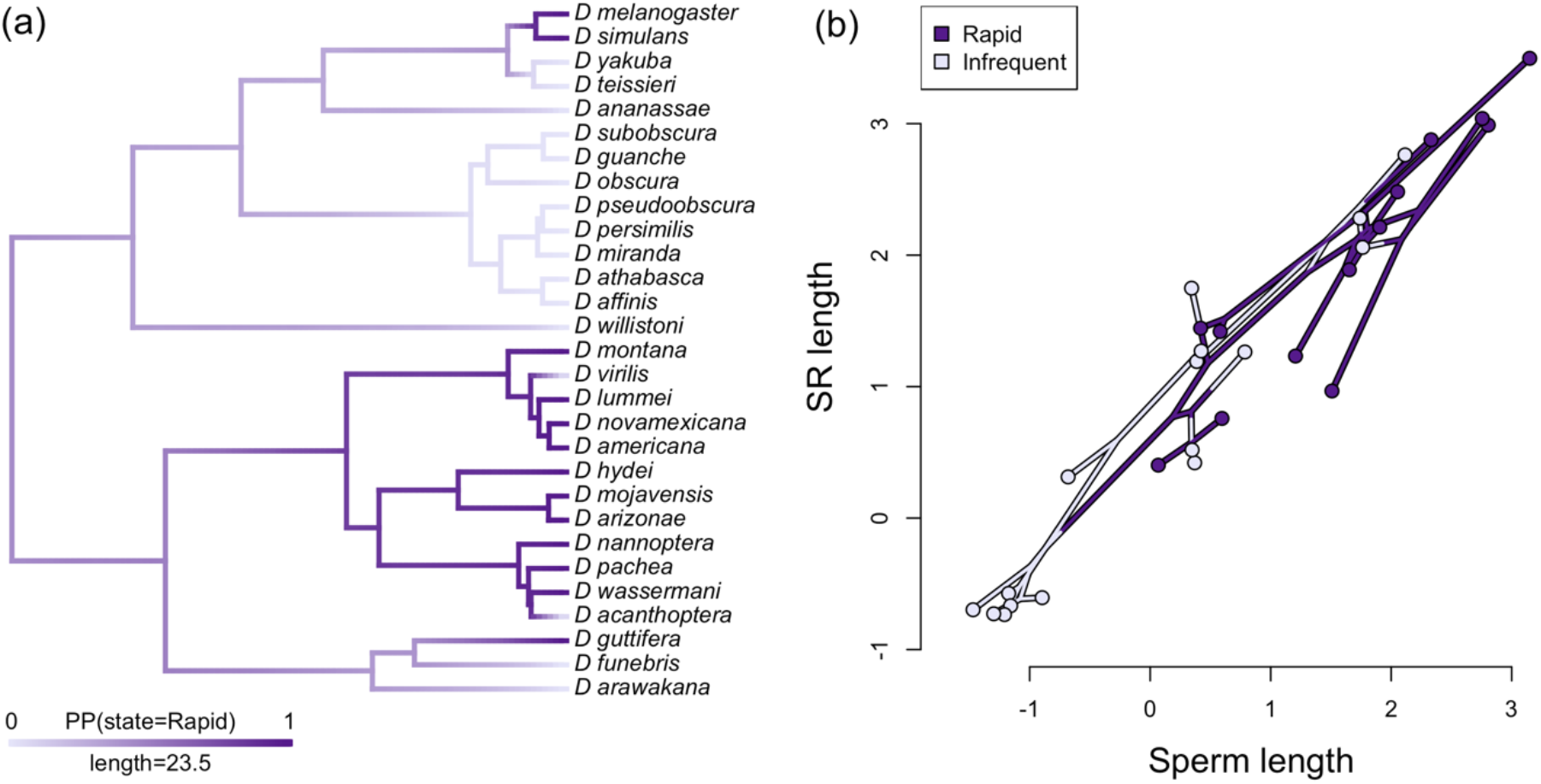
(a) Evolutionary reconstruction of variation in remating rate across the *Drosophila* phylogeny, shown as a posterior density map from 1000 stochastic mappings of remating rate on the tree. Tips represent observed trait values. Dark purple signifies that more than half of females remated within 24 hours (rapid remating), while light purple females remated less frequently (infrequent remating). This tree includes all species shown in Figure 1 except *D. immigrans* which does not have remating data. (b) Phylomorphospace for the relationship between sperm length and SR length for *Drosophila* species in either infrequent or rapid remating groups. The color of each node represents the character state from a single stochastic mapping of remating rate on the phylogeny. SR length and sperm length data is log- transformed.

Using the same 15-species dataset, we also tested for associations between MRT mass and our other traits of interest, because MRT mass has also been used as a proxy for the strength of postmating sexual selection (e.g., Holman et al. 2008). We found that MRT mass significantly predicted female remating rate (though this association is not significant in the opposite direction; Table S7); specifically, species where males had heavy reproductive tracts had females that remated quickly (Table S7). This result is consistent with Holman et al. (2008), who detected the same association in the *D. obscura* group. However, MRT mass is not consistently associated with any other aspects of reproductive morphology (SR length, ST area, ST pigmentation, FRT mass, testis length, testis color, or sperm length).

Using our expanded dataset, where species are assigned to two remating states—rapid or infrequent remating groups—we tested this question across 29 species for three of our post- mating traits (SR length, testis length, or sperm length). We found no significant correlation between female remating rate and any of these three measures of female or male reproductive morphology, when taking phylogeny into account (Figure S8, Figure S9).

Finally, using our expanded dataset of 29 species, we fit a multivariate Brownian model to test whether rates of SR or sperm length evolution (or their coevolution) differed on branches of the phylogeny where female remating is reconstructed as either rapid or infrequent (as determined by stochastic mapping of ancestral states for this discrete trait; see above and Figure 4). The model with the lowest AIC value was one which allowed different rates of evolution for SR length (σ^2^ = 0.084 in rapidly remating groups vs. σ^2^ = 0.055 in infrequently remating groups) but not for sperm length (σ^2^ = 0.059) (i.e. Model 2c in Table S10). However, neither this model nor any other model that incorporated variable rates or rate covariation in rapidly versus infrequently remating groups, fit significantly better than a model with a single rate of evolution for both groups (Model 1, Table S10), based on a comparison of AIC scores (Burnham and Anderson 2004). This included models that allowed the evolutionary covariance between SR and sperm length to differ between rapidly and infrequently remating groups (Table S10), as might be expected if rapid remating consistently elevated the opportunity for stronger male-female postmating coevolution. Moreover, the correlation between sperm length and SR length in infrequently mating lineages was r = 0.945, which is slightly (but not significantly) higher than the correlation in rapidly remating lineages (r = 0.903; Figure 4b). This indicates that the correlation between sperm and the SR is strong across *Drosophila* species, regardless of remating rate. Overall, we found no evidence that rapid remating was consistently associated with the exaggerated size or evolutionary rate of either SR length in females or sperm or testis length in males.

### 3.5 Is remating rate more closely associated with postmating trait evolution than premating or perimating trait evolution?

Among all our traits (Table S7, Figure S8, S9), the only significant phylogenetically corrected association we detected with female remating rate was a significant negative association between mating duration and remating rate (mean r = -0.512) in our expanded (n=29 species) dataset (Table S7, Figure S10). This suggests that rather than being consistently associated with postmating traits, female remating rate appears to be more closely associated with a perimating trait.

We also tested whether life history—specifically age to reproductive maturity—might shape reproductive trait evolution. We found no evidence that the age of female reproductive maturity is associated with female remating rate, female postmating reproductive morphology, premating reproductive behavior, or perimating behavior in either our original 15 species (Table S7) or our expanded dataset (Table S8).

## 4 Discussion

Sexual selection is proposed to be a powerful driver of trait diversity, including in reproductive traits that act during and after mating. In this study, we used a comparative phylogenetic framework to assess the rate and pattern of evolution of both male and female pre- and postmating traits across up to 30 *Drosophila* species, and to examine evidence for the role of sexual selection in shaping their evolution. We found that reproductive traits generally evolve more rapidly than non-reproductive traits (prediction 1). We found no evidence that male postmating reproductive traits are evolving faster than female traits (prediction 2), but did find clear evidence of postmating trait coevolution between males and females (prediction 3).

Interestingly, rapidly and infrequently remating lineages did not differ in their rates of postmating trait evolution, or in the strength of association between sperm and sperm storage traits (prediction 4). Instead, we found that female remating rate evolution was more closely associated with perimating than postmating trait evolution (prediction 5), specifically via an association between remating rate and mating duration. As we expand upon below, these results suggest that the evolution of our postmating traits is primarily driven by intersexual and female-mediated selection, with a complex and likely secondary role for remating rate in shaping this evolution. Our findings also highlight that the targets of postmating selection can be multimodal, and can vary over evolutionary time in response to other factors that might shape or constrain their evolution.

### 4.1 Intersexual and female-mediated selection drives the evolution of our postmating traits

We initially predicted that male traits might evolve more rapidly because they are expected to experience multiple forms of sexual selection (both intra- and intersexual), while female postmating traits are only subject to intersexual selection. Similar logic has been used to explain the exaggeration of traits that mediate premating interactions (Clutton-Brock 1983).

However, we found no evidence that the evolutionary rates of male postmating traits generally outpace female traits. This finding may underscore a fundamental difference between premating and postmating stages of reproductive interaction: Male competition and female choice often occur separately in a premating context; however, this distinction becomes obscured after mating, where competition and choice occur concurrently inside the female reproductive tract (Lüpold et al. 2020, Garlovsky et al. 2023). The temporal and physical co-occurence of postmating interactions amplifies the opportunity for direct cross-talk between male- and female-mediated processes, and thereby between intra- and intersexual selection. It also amplifies the potential role of female-mediated selection, as the female herself is the biological arena within which these interactions play out.

Our findings are consistent with both this sustained cross-talk between male- and female-mediated evolution, and an amplified role for females. Specifically, strong phylogenetically independent covariation between male (sperm length) and female (SR length) traits (Figure 4) indicates a significant role for male-female coevolution in shaping these traits across *Drosophila* species, as has also been inferred in other studies in *Drosophila* (Pitnick et al. 1999; Holman et al. 2008) and other animal systems (e.g., Briskie et al. 1997, Presgraves et al. 1999, Minder et al. 2005, Anderson et al. 2006, Beese et al. 2009). Moreover, our findings suggest that female-mediated processes are as influential as male-mediated processes in driving the evolution of these reproductive traits. A component of Eberhard’s (1996) proposal that female choice could drive sexual trait evolution was that greater female reproductive tract complexity could be an effective mechanism of exercising choosiness over sperm. Others have since inferred that increasing SR length might be the initial microevolutionary trigger (e.g. Miller and Pitnick 2002) and the sustained macroevolutionary driver (e.g., Lüpold et al. 2016, Higginson et al. 2012) of male testis and/or sperm length exaggeration. Whether females are the primary initiators of postmating trait evolution is challenging to disentangle in our study. However, several of our observations fit with Eberhard’s conception of evolution initiated and propelled by cryptic female choice. In particular, instead of more rapid male evolution, female traits either match or outpace male trait evolution. Indeed, two female traits—FRT mass and ST area—have the fastest estimated rates of evolutionary change among all morphological postmating traits we measured (Table S7). Moreover, within our pattern of tight co-evolution between testis or sperm length and sperm storage organ morphology, the magnitude of SR morphological change either matches or exceeds the magnitude of male morphological change (e.g., the relationship between standardized trait values is greater than one; Figure 2). A similar ‘positive allometric’ relationship was previously noted by Pitnick et al. (1999), who observed an even greater proportional change in SR length compared to sperm length evolution among *Drosophila* species.

### 4.2 Postmating morphological trait evolution is decoupled from remating rate

A second major observation in our study is that patterns of postmating morphological evolution are decoupled from our measure of the expected intensity of postmating sexual selection: female remating rate. Female remating rate is a frequent proxy for the strength of postmating sexual selection because the opportunity for both sperm competition and cryptic female choice is expected to be greater when different males’ sperm overlap in a female’s reproductive tract (Markow 2002). However, once phylogenetic relationships are taken into account, we do not find any sustained association between female remating rate and the exaggeration of most reproductive morphology (SR length, ST area, ST color, testis length, testis color, sperm length, or FRT mass) or the rate of evolution of these morphological traits.

Interestingly, using a similar phylogenetically corrected comparison, Holman et al. (2008) also did not detect a relationship between remating rate and SR or sperm length evolution within the *Drosophila obscura* species group—a clade that has sperm heteromorphism. Our findings indicate that this observation extends to both sperm-heteromorphic and sperm-monomorphic groups of *Drosophila*.

Several factors could contribute to this intriguing macroevolutionary decoupling of remating rate and postmating trait evolution. First, it is possible that the intensity of post-copulatory competition does not simply or directly translate to the evolutionary response to this selection, even if female preference drives male-female morphological coevolution. This inference is suggested by recent theory, which found that cryptic female choice can drive male-female coevolution in reproductive traits even when the strength of cryptic female choice is weak (Kustra and Alonzo 2023). Consistent with this, we found that the correlation between sperm and SR length was identical, regardless of variation in remating rate (i.e., in both rapid and infrequent remating clades; Figure 4). Moreover, in lineages that have more recently transitioned to infrequent remating, we find no evidence of a reduction in sperm or SR length (Figure 1, Figure 4), suggesting that the forces acting on these traits were sustained regardless of the specific current remating rate.

A second, non-exclusive explanation for the decoupling of remating rate and morphological trait evolution appeals to the fact that remating rate can be both a response to, and a driver of, postmating sexual selection. Alternating between these roles across the evolutionary history of *Drosophila* could obscure or erase a pattern of coevolution with postmating morphological evolution. This explanation is consistent with elements of our ancestral state reconstructions for remating rate (Figure 4) and for SR and sperm length (Figure 1). These reconstructions indicate that the major transition to exaggerated SR and sperm length occurred early (∼34 MYA) in the evolutionary history of subgenus *Drosophila* (at the base of this clade, in Figure 1), whereas the major transition from infrequent to rapid remating appears to occur later (∼24 MYA) within that clade (on the branch giving rise to the subgroup containing *D. montana* through *D. acanthoptera*, Figure 4). If the exaggeration of both SR and sperm length preceded the switch to rapid remating in one of our major clades, rapid remating rate could not have driven this trait elaboration. These major consecutive transitions also help explain why remating rate is statistically associated with SR and sperm length among our contemporary species, but does not remain so after phylogenetic correction. Unreplicated state shifts (“Darwin’s scenario”; Maddison and FitzJohn 2015) can mislead inferences when correlations between traits are due to one or a few major transitions in the history of each trait (Uyeda et al. 2018), underscoring the value of analyzing these relationships within a comparative phylogenetic framework.

A dynamic evolutionary role for remating rate—as both an agent and an outcome of postmating sexual selection—is supported by several other elements of our analysis, as well as many mechanistic studies of this trait (e.g., Alonzo and Pizzari 2013, Laturney et al. 2018). Our estimated evolutionary rate for remating—and for other related behavioral traits—is higher than for most of our morphological postmating traits (Table S5), suggesting greater evolutionary lability and more localized clade-specific responses to selection in these traits. Moreover, in species with short copulation durations, females tend to remate more rapidly—a phylogenetically independent trait correlation that suggests coordinated evolution of specific mating strategies that shape the relationship between mating and postmating outcomes. For instance, shortened copulation duration in some species could be a female-mediated mechanism to increase remating rate, via its modulation of female exposure to male seminal fluid proteins. These proteins— transferred during copulation, often after the primary transfer of sperm (Gilchrist and Partridge 2000; see also Wigby et al. 2009)—play an important role in postmating responses, including suppressing female remating (Chapman and Davies 2004) (reviewed in Pitnick et al. 2009).

Short matings may attenuate the transfer of seminal fluid proteins, thereby reducing the period that females remain refractory between matings. Conversely, rapid remating after short copulations might function to replenish depleted sperm stores (i.e. to counter sperm limitation). Both scenarios are consistent with remating rate (and associated mating traits) being a dynamic response to the persistent tension between male and female interests over determining the paternity outcome of matings (Arnqvist et al. 2000, Härdling and Kaitala 2005). These possibilities also allow for the relationship between remating rate and the intensity of postmating sexual selection to change over evolutionary time, as this female behavioral trait varies in its roles as a driver and a target of postmating selection. This varied role is more complex than the widespread expectation (and in some cases inference; Markow 2002) that remating rate plays a strong consistent role in imposing postmating sexual selection.

### 4.3 The evolutionary targets and responses to postmating sexual interactions change over time

The above inferences indicate that targets and drivers of postmating sexual selection can shift over time. Beyond dynamic coevolutionary interactions, our data suggest these shifts might also be shaped by the emergence of mechanical or physiological limits on continued responses to selection. For instance, we infer rapid morphological elaboration of both SR and sperm length early in the history of the *Drosophila* subgenus (Figure 1), consistent with experimental evolution that demonstrates that these traits can rapidly respond to postmating selection under relatively unconstrained lab conditions (Miller and Pitnick 2002). However, many extant species in this clade now have very large proportional investments in reproductive tissues. For example, in *D. mojavensis* reproductive tissues make up 20.8% and 27.6% of total body mass, in males and females respectively (Table S2, Table S3). In these lineages, there may be substantially less scope for continued increases in trait size or elaboration in response to postmating sexual selection, even under conditions of rapid remating. Upper limits on reproductive organ size or investment in these clades could explain why traits like sperm, SR, and testis length have relatively modest overall rates of evolution compared to several other (especially behavioral) reproductive traits (Table S5), when estimated across the entire 47 MY timescale assessed here. They also further explain why macroevolutionary change in these internal morphological characters can become decoupled from more dynamic traits like remating rate.

In clades where these emerging constraints limit further evolution, other traits might instead be responding to ongoing postmating sexual selection. Apart from the peri- and post-mating behavioral traits that we observe to be relatively rapidly evolving, these other potential targets include additional postmating traits not assessed here, such as the novel sperm storage organ observed in *D. nigricruria* (Pitnick et al. 1999) (a species not included here), or seminal fluid protein identity. Seminal fluid protein genes in particular have been shown to be under strong positive selection and rapidly evolving in *Drosophila* males (Wagstaff and Begun 2005, Haerty et al. 2007), consistent with ongoing intense sexual selection acting on these genes.

Moreover, evolution of these loci interacts with traits like remating rate (Sirot et al. 2015). For example, the seminal fluid protein sex peptide—which suppresses female remating—appears to evolve rapidly on the evolutionary branch leading to *D. ananassae* (McGeary and Findlay 2020)—a species for which we observe an extremely infrequent remating rate. If these proteins are the focus of selection imposed by remating frequency, for example, their molecular evolutionary rates should be significantly associated with quantitative or qualitative remating rate variation among our species, as we originally predicted for morphological traits. This testable association might be particularly evident in lineages whose postmating morphologies are already highly exaggerated and therefore constrained in terms of further evolutionary responses.

### 4.4 What drives the diversification of postmating traits?

Our goal here was to address a set of predictions about the evolution of postmating reproductive traits among up to 30 *Drosophila* species across a timescale of 47 MY. Among the remarkable macroevolutionary diversity we observe, we find that the pace of female trait evolution is matching or possibly exceeding that of male traits, consistent with roles for female-mediated and male-female coevolution in driving these traits. In contrast, remating rate does not consistently explain the rate or pattern of postmating trait evolution, including the initial exaggeration of these traits in groups where they are highly elaborated. Our analyses indicate that the factors shaping postmating reproductive trait evolution are not always straightforward. Many traits could act as either drivers of or responses to postmating selection over their evolutionary history; this includes remating rate itself, which complicates its role as a simple proxy for the strength of postmating sexual selection. Moreover, factors like the emergence of upper limits on some trait responses, mean that both the mechanism and target of postmating selection can change over time. These dynamics are a double-edged sword: while they make it complex to interpret causal relationships among contemporary traits, they likely also amplified the diversification of postmating traits during their divergence history, producing the richness of reproductive traits, trait values, and trait combinations we observe today.

## Data availability statement

The data and code underlying this article are available in the Dryad Digital Repository, at https://dx.doi.org/[doi].

## Acknowledgments

We thank Stuart J. Macdonald for providing the *D. melanogaster* line used in this study, Brandon S. Cooper for providing *D. yakuba* and *D. teissieri,* Dean M. Castillo for collecting *D. pseudoobscura* and *D. persimilis,* and Scott Hawley for providing *D. ananassae, D. simulans,* and *D. mojavensis*. We also thank the Cornell *Drosophila* Species Stock Center for providing the rest of the lines used in this study. We thank Spencer R. Hall for the use of his microbalance. We thank Matthew W. Hahn, Kimberly A. Rosvall, and Michael J. Wade for their helpful comments on the manuscript. The research was supported by Indiana University Department of Biology funding to LCM and BP, as well as the NIH Common Themes in Reproductive Diversity (CTRD) training grant funding to BP.

