## Supplementary Material for "Females drive postmating reproductive trait evolution across *Drosophila* species, but not via remating rate"

**Supplementary Methods**

*2.2 Developmental and behavioral data collection*

*2.2.1 Age of female reproductive maturity*

To accurately estimate female mating and remating traits, we first had to experimentally determine the female age of reproductive maturity for each species. We did this using a previously established experimental approach (e.g., Pitnick et al. 1995, Snook and Markow 2001, Holman et al. 2008). Specifically, we set up five replicate crosses, each of which paired an individual 3-day-old female with two reproductively mature males (either 7 or 14 days old, depending on the species). These crosses were set up between 9-11 am, within two hours of the lights coming on. If at least 80% of females mated within 2 hours of observation, we dissected the reproductive tract of each female to confirm the transfer of sperm. If there was no sperm transferred, we set up new crosses with older males. If sperm was present, we next used 2-day-old females, to assess whether at least 80% of females mated within 2 hours and received sperm. This process of reducing female age was continued until fewer than 80% of females mated. (Note that no species had an age of female reproductive maturity <2 days; see Results.) If for a specific line, 80% of 3-day-old females did not mate, we increased female age by next setting up 4-day-old females for this line, and so forth until we identified the day on which the majority of virgin females would mate. The youngest age at which at least 80% of females mated was considered the age of reproductive maturity for that species. Once that age was determined, we set up an additional five replicates to confirm that 80% of females still mate in the determined period (for a total replication of n = 10). Crosses were completed in blocks with 3-4 randomly chosen species at a time. Mating latency and mating duration were also recorded for each mating.

*2.2.2 Female remating rate*

We quantitatively estimated the female remating rate for each line/species by experimentally determining the minimum interval within which the majority of previously mated females would remate (female intermating interval), based on established methods (e.g., Snook and Markow 2001, Holman et al. 2008). Specifically, starting with females at their previously determined age of sexual maturity, we set up crosses in which individual virgin females were paired with two virgin, reproductively mature males (males were either 7 or 14 days old, depending on the species). These crosses were set up between 9-11 am, within two hours of the lights coming on. Once females mated, males were removed from the vial. Females were left in isolation until the following morning, when they were set up with two new reproductively mature males and observed for two hours. This process was repeated daily until the female remated. We identified the female intermating interval for at least five females of each species. We set up four crosses per species, in blocks with 3-4 randomly chosen species at a time. Mating latency and mating duration were also recorded for both first and second matings.

For species in which females always remated within 48 hours (*D. simulans, D. americana, D. novamexicana, D. lummei, D. mojavensis,* and *D. hydei)*, we repeated the assessment with an additional remating opportunity four hours after the end of the first mating opportunity (in the afternoon of the first day, from 3-5pm). Thus, we could assess whether these species would remate multiple times in one day. We again identified the female intermating interval for at least five females of each species. We ultimately transformed this variable into the female remating rate by taking the inverse of the intermating interval.

*2.3 Morphological data collection*

*2.3.1 Thorax length, reproductive tract length, and pigmentation*

We collected both female and male virgins and aged them to reproductive maturity (as previously determined experimentally). We measured thorax length for all flies; this is a standard measure of body size in studies of *Drosophila* because it strongly correlates with the size of other characters such as wing length (Robertson and Reeve 1952). We then dissected the reproductive tract into Grace's Insect Medium. For females, we separated the seminal receptacle (SR) and a randomly chosen spermatheca (ST) from the rest of the reproductive tract and transferred them to a slide. For males, a randomly chosen testis was separated from the rest of the reproductive tract and transferred to a slide. In both cases, we then applied a cover slip and took images under an EVOS FL digital inverted microscope. These images were later analyzed in ImageJ to measure the length of the SR, the height and width of the ST, and the length of the testis.

These images were also used to assess pigmentation intensity of the ST and the testis in ImageJ. The function of male and female postmating reproductive trait pigmentation is unknown in *Drosophila*, but we included these traits because they clearly differed between species. To do this, ST images were converted to 8-bit grayscale, and a 15 μm diameter circle within the ST was assessed for mean grayvalue. Testes were also classified into discrete color categories based on visual assessment (see Figure S1 for examples of each color). Length and pigmentation measurements were replicated across five individuals per species.

*2.3.2 Body mass and reproductive tract mass*

We collected both female and male virgins and aged them to reproductive maturity. We measured body and reproductive tract mass according to Holman et al. (2008). Specifically, flies were dissected in distilled water onto a piece of preweighed foil. The female reproductive tract (FRT), which included the bursa, SR, ST, parovaria, and ovaries, was removed and transferred to a second piece of pre-weighed foil. The male reproductive tract (MRT), which included testes, seminal vesicles, accessory glands, and the ejaculatory bulb, was removed and transferred to a second piece of pre-weighed foil. For both males and females, each foil (holding either the body or the reproductive system) was then placed in a drying oven at 60°C for 18 ± 1 h and weighed on a Mettler Toledo UMX2 balance to the nearest 0.1 microgram. Mass measurements were replicated across five individuals per species.

*2.3.3 Sperm length*

Sperm length was also experimentally assessed for a subset of ten of the species for which other morphological traits were directly measured (specifically, *D. melanogaster, D. simulans, D. yakuba, D. teissieri, D. ananassae, D. pseudoobscura, D. persimilis, D. mojavensis, D. arawakana,* and *D. immigrans*). The five experimental species with longest sperm (*D. virilis, D. lummei, D. americana, D. novamexicana,* and *D. hydei*) were not assessed, as this experimental procedure is not effective for accurately measuring sperm longer than ~2 mm. We dissected sperm out of the seminal vesicles of reproductively mature virgin males into Grace's Insect Medium on a slide. Sperm was gently teased apart with a probe when necessary. The slides were stained with SpermBlue (Microptic), then we took photos under an EVOS FL digital inverted microscope. These photos were later analyzed in ImageJ to measure sperm length for 5-10 randomly chosen sperm per individual. We repeated these measurements across five individuals per species.

**Supplementary Tables and Figures**

**Table S1.** List of fly lines used and their sources.

| Species | Line | Source |
|---|---|---|
| ***D. melanogaster*** | B4 (California) | Drosophila Synthetic Population Resource |
| ***D. simulans*** | 14021-0251.006 | Cornell Drosophila Species Stock Center |
| ***D. yakuba*** | yak_S09_L38 | Sao Tome, 2009 (Cooper et al. 2017) |
| ***D. teissieri*** | tei_B09_L4 | Bioko, 2009 (Cooper et al. 2017) |
| ***D. ananassae*** | 14024-0371.13 | Cornell Drosophila Species Stock Center |
| ***D. pseudoobscura*** | LCMS2-5 | Lamoille Canyon, NV (Castillo and Moyle 2019) |
| ***D. persimilis*** | CR1 | Sierra, CA (Castillo and Moyle 2019) |
| ***D. willistoni*** | 14030-0811.17 | Cornell Drosophila Species Stock Center |
| ***D. americana*** | 15010-1041.29 | Cornell Drosophila Species Stock Center |
| ***D. novamexicana*** | 15010-1031.08 | Cornell Drosophila Species Stock Center |
| ***D. lummei*** | 15010–1011.07 | Cornell Drosophila Species Stock Center |
| ***D. virilis*** | Genome line | Cornell Drosophila Species Stock Center |
| ***D. mojavensis*** | 15081-1352.22 | Cornell Drosophila Species Stock Center |
| ***D. hydei*** | 15085-1641.76 | Cornell Drosophila Species Stock Center |
| ***D. arawakana*** | 15182-2261.03 | Cornell Drosophila Species Stock Center |
| ***D. immigrans*** | 15111-1731.12 | Cornell Drosophila Species Stock Center |

**Table S2.** Female reproductive data across 15 *Drosophila* species. Data are represented as mean values ± standard deviations where applicable. ARM is age of reproductive maturity. n = 5 for all morphological traits, and n = 5-10 for behavioral traits.

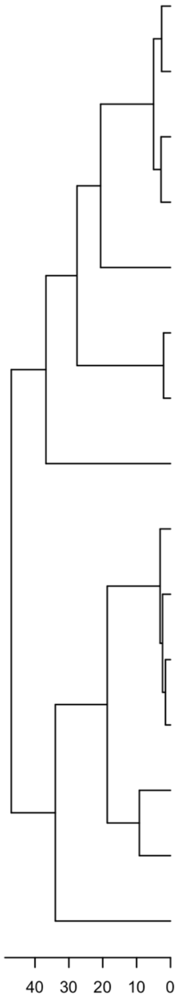

| Species | ARM (days) | Remating interval (days) | Mating latency (min) | Mating duration (min) | Thorax length (mm) | SR length (mm) | ST area ($mm^2$) | ST pigment intensity (1/grayvalue) | Non-RT body mass (mg) | RT mass (mg) |
|---|---|---|---|---|---|---|---|---|---|---|
| *D. melanogaster* | 2 | 2.57 ± 2.64 | 7.3 ± 3.9 | 17.9 ± 2.8 | 0.882 ± 0.040 | 2.134 ± 0.172 | 0.005 ± 0.000 | 0.013 ± 0.001 | 0.288 ± 0.015 | 0.095 ± 0.019 |
| *D. simulans* | 2 | 1.10 ± 0.55 | 6.4 ± 4.7 | 16.0 ± 3.9 | 0.821 ± 0.069 | 1.500 ± 0.104 | 0.003 ± 0.001 | 0.017 ± 0.002 | 0.229 ± 0.038 | 0.072 ± 0.032 |
| *D. yakuba* | 7 | 6.80 ± 4.49 | 14.4 ± 4.9 | 38.0 ± 5.1 | 0.794 ± 0.064 | 1.679 ± 0.243 | 0.005 ± 0.001 | 0.012 ± 0.001 | 0.232 ± 0.036 | 0.116 ± 0.014 |
| *D. teissieri* | 3 | 4.50 ± 2.88 | 22.2 ± 23.0 | 35.0 ± 6.4 | 0.869 ± 0.086 | 1.523 ± 0.059 | 0.004 ± 0.001 | 0.013 ± 0.001 | 0.236 ± 0.037 | 0.091 ± 0.025 |
| *D. ananassae* | 4 | 23.33 ± 5.72 | 11.2 ± 8.2 | 5.3 ± 0.8 | 0.935 ± 0.042 | 3.537 ± 0.371 | 0.007 ± 0.001 | 0.008 ± 0.000 | 0.378 ± 0.062 | 0.050 ± 0.020 |
| *D. pseudoobscura* | 3 | 14.60 ± 9.40 | 4.2 ± 4.6 | 9.2 ± 0.8 | 1.017 ± 0.073 | 0.480 ± 0.066 | 0.005 ± 0.001 | 0.014 ± 0.001 | 0.348 ± 0.027 | 0.082 ± 0.033 |
| *D. persimilis* | 3 | 4.67 ± 4.63 | 24.7 ± 34.9 | 17.8 ± 4.7 | 1.048 ± 0.048 | 0.513 ± 0.028 | 0.005 ± 0.002 | 0.022 ± 0.001 | 0.324 ± 0.018 | 0.206 ± 0.035 |
| *D. willistoni* | 4 | 5.86 ± 2.27 | 6.3 ± 4.7 | 14.9 ± 2.1 | 0.777 ± 0.052 | 5.747 ± 0.362 | 0.008 ± 0.001 | 0.008 ± 0.000 | 0.221 ± 0.027 | 0.056 ± 0.014 |
| *D. virilis* | 14 | 2.40 ± 0.55 | 65.8 ± 39.0 | 4.8 ± 0.8 | 1.190 ± 0.068 | 9.788 ± 0.068 | 0.018 ± 0.002 | 0.011 ± 0.001 | 0.660 ± 0.089 | 0.144 ± 0.043 |
| *D. lummei* | 9 | 0.90 ± 0.65 | 37.6 ± 16.5 | 3.6 ± 0.9 | 1.283 ± 0.043 | 11.962 ± 0.365 | 0.013 ± 0.002 | 0.011 ± 0.000 | 0.699 ± 0.082 | 0.089 ± 0.025 |
| *D. novamexicana* | 5 | 0.60 ± 0.22 | 31.8 ± 24.4 | 3.2 ± 0.4 | 1.196 ± 0.045 | 9.143 ± 0.995 | 0.017 ± 0.002 | 0.009 ± 0.000 | 0.899 ± 0.068 | 0.114 ± 0.011 |
| *D. americana* | 5 | 0.80 ± 0.27 | 35.0 ± 29.5 | 3.4 ± 0.5 | 1.298 ± 0.066 | 6.610 ± 0.769 | 0.017 ± 0.001 | 0.012 ± 0.001 | 0.721 ± 0.060 | 0.129 ± 0.012 |
| *D. hydei* | 6 | 0.50 ± 0.00 | 11.2 ± 5.3 | 2.2 ± 0.4 | 1.189 ± 0.046 | 33.014 ± 2.871 | 0.016 ± 0.001 | 0.008 ± 0.000 | 0.832 ± 0.098 | 0.098 ± 0.024 |
| *D. mojavensis* | 8 | 1.00 ± 0.61 | 6.6 ± 3.6 | 3.4 ± 1.1 | 0.841 ± 0.018 | 4.133 ± 0.542 | 0.001 ± 0.000 | 0.006 ± 0.000 | 0.216 ± 0.063 | 0.092 ± 0.059 |
| *D. arawakana* | 5 | 17.00 ± 3.74 | 13.6 ± 10.0 | 30.4 ± 5.2 | 0.889 ± 0.044 | 3.291 ± 0.309 | 0.008 ± 0.001 | 0.011 ± 0.001 | 0.239 ± 0.050 | 0.062 ± 0.019 |

**Table S3.** Male reproductive data across 15 *Drosophila* species. Data are represented as mean values ± standard deviations where applicable. ARM is age of reproductive maturity (note that males were only assessed for reproductive maturity at 7 and 14 days of age). n = 5 for all morphological traits.

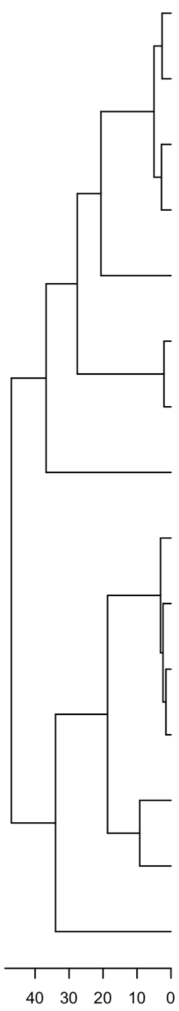

| Species | ARM (days) | Thorax length (mm) | Testis length (mm) | Testis color | Sperm length (mm) | Non-RT Body mass (mg) | MRT mass (mg) |
|---|---|---|---|---|---|---|---|
| *D. melanogaster* | 7 | 0.789 ± 0.043 | 2.858 ± 0.130 | Yellow | 1.814 ± 0.033 | 0.229 ± 0.027 | 0.025 ± 0.004 |
| *D. simulans* | 7 | 0.714 ± 0.024 | 1.660 ± 0.109 | Yellow | 1.069 ± 0.018 | 0.160 ± 0.020 | 0.028 ± 0.002 |
| *D. yakuba* | 7 | 0.683 ± 0.048 | 2.105 ± 0.169 | Yellow | 1.416 ± 0.029 | 0.154 ± 0.009 | 0.024 ± 0.004 |
| *D. teissieri* | 7 | 0.773 ± 0.029 | 2.093 ± 0.184 | Yellow | 1.446 ± 0.063 | 0.149 ± 0.011 | 0.021 ± 0.003 |
| *D. ananassae* | 7 | 0.829 ± 0.050 | 3.252 ± 0.310 | Yellow | 2.194 ± 0.041 | 0.219 ± 0.020 | 0.023 ± 0.004 |
| *D. pseudoobscura* | 7 | 0.891 ± 0.052 | 0.911 ± 0.035 | Orange | 0.298 ± 0.003 | 0.180 ± 0.023 | 0.030 ± 0.005 |
| *D. persimilis* | 7 | 0.898 ± 0.038 | 0.840 ± 0.013 | Orange | 0.314 ± 0.004 | 0.280 ± 0.012 | 0.038 ± 0.009 |
| *D. willistoni* | 7 | 0.670 ± 0.055 | 2.055 ± 0.139 | Yellow | 1.410 ± 0.040 | 0.162 ± 0.013 | 0.031 ± 0.006 |
| *D. virilis* | 14 | 1.178 ± 0.022 | 8.052 ± 0.178 | Orange | -- | 0.485 ± 0.025 | 0.057 ± 0.008 |
| *D. lummei* | 14 | 1.162 ± 0.064 | 9.800 ± 0.478 | Orange | -- | 0.505 ± 0.026 | 0.052 ± 0.004 |
| *D. novamexicana* | 7 | 1.203 ± 0.035 | 8.081 ± 0.564 | Orange | -- | 0.730 ± 0.101 | 0.044 ± 0.009 |
| *D. americana* | 7 | 1.256 ± 0.073 | 7.887 ± 0.517 | Orange | -- | 0.573 ± 0.065 | 0.052 ± 0.007 |
| *D. hydei* | 7 | 1.231 ± 0.113 | 25.762 ± 2.086 | Yellow | -- | 0.370 ± 0.015 | 0.062 ± 0.010 |
| *D. mojavensis* | 14 | 0.820 ± 0.039 | 2.522 ± 0.164 | Yellow | 1.788 ± 0.028 | 0.185 ± 0.014 | 0.048 ± 0.005 |
| *D. arawakana* | 14 | 0.798 ± 0.055 | 2.408 ± 0.424 | White | 1.471 ± 0.149 | 0.155 ± 0.023 | 0.011 ± 0.003 |

**Table S4.** Mean trait values for the expanded dataset. Footnotes indicate sources for individual data points. Any data without a footnote is from this paper.

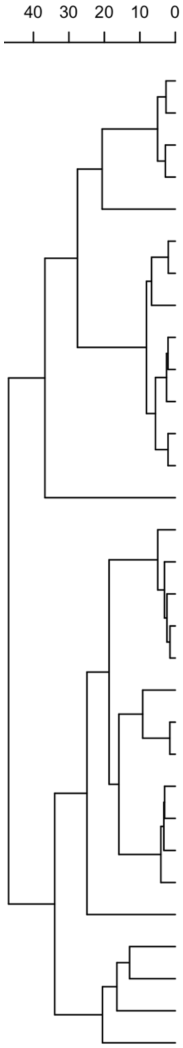

| | Remating rate | Female ARM (days) | Mating duration (min) | Female thorax length (mm) | SR length (mm) | Male thorax length (mm) | Testis length (mm) | Sperm length (mm) |
|---|---|---|---|---|---|---|---|---|
| *D. melanogaster* | Rapid | 2 | 17.9 | 0.882 | 2.134 | 0.789 | 2.858 | 1.814 |
| *D. simulans* | Rapid | 2 | 16.0 | 0.821 | 1.496 | 0.714 | 1.660 | 1.069 |
| *D. yakuba* | Infrequent | 7 | 38.0 | 0.794 | 1.679 | 0.683 | 2.105 | 1.416 |
| *D. teissieri* | Infrequent | 3 | 35.0 | 0.869 | 1.523 | 0.773 | 2.093 | 1.446 |
| *D. ananassae* | Infrequent | 4 | 5.3 | 0.935 | 3.537 | 0.829 | 3.252 | 2.194 |
| *D. subobscura* | Infrequent[1] | 6[1] | 4.7[1] | 1.158[1] | 0.546[1] | 1.085[2] | -- | 0.408[1] |
| *D. guanche* | Infrequent[1] | 6[1] | 8.5[1] | 1.071[1] | 0.483[1] | 0.992[2] | -- | 0.273[1] |
| *D. obscura* | Infrequent[1] | 4[1] | 8.6[1] | 1.197[1] | 0.498[1] | 1.127[2] | -- | 0.230[1] |
| *D. persimilis* | Infrequent | 3 | 17.8 | 1.048 | 0.513 | 0.898 | 0.840 | 0.314 |
| *D. pseudoobscura* | Infrequent | 3 | 9.2 | 1.017 | 0.480 | 0.891 | 0.911 | 0.298 |
| *D. miranda* | Infrequent[1] | 5[1] | 7.6[1] | 1.250[1] | 0.564[1] | 1.185[2] | -- | 0.309[1] |
| *D. athabasca* | Infrequent[1] | 6[1] | 9.6[1] | 1.033[1] | 3.564[1] | 0.951[2] | -- | 1.527[1] |
| *D. affinis* | Infrequent[1] | 5[1] | 1.6[1] | 1.080[1] | 1.368[1] | 0.920[2] | -- | 0.506[1] |
| *D. willistoni* | Infrequent | 4 | 14.9 | 0.777 | 5.747 | 0.670 | 2.055 | 1.410 |
| *D. montana* | Rapid[3] | 4[4] | 4.3[5] | 1.48[4] | 3.43[6] | 1.41[4] | -- | 3.34[4] |
| *D. virilis* | Infrequent | 14 | 4.8 | 1.190 | 9.788 | 1.178 | 8.052 | 5.70[4] |
| *D. lummei* | Rapid | 9 | 3.6 | 1.283 | 11.962 | 1.162 | 9.800 | 7.79[4] |
| *D. novamexicana* | Rapid | 5 | 3.2 | 1.196 | 9.143 | 1.203 | 8.081 | 6.72[4] |
| *D. americana* | Rapid | 5 | 3.4 | 1.298 | 6.610 | 1.256 | 7.887 | 5.22[4] |
| *D. hydei* | Rapid | 6 | 2.2 | 1.189 | 33.014 | 1.231 | 25.762 | 23.32[4] |
| *D. mojavensis* | Rapid | 8 | 3.4 | 0.841 | 4.133 | 0.820 | 2.522 | 1.788 |
| *D. arizonae* | Rapid[3] | 3[4] | 1.3[5] | 1.03[4] | 4.24[6] | 0.95[4] | -- | 1.52[4] |
| *D. wassermani* | Rapid[3] | 4[4] | 14.2[5] | 1.16[4] | 2.63[6] | 1.07[4] | 7.76[7] | 4.52[4] |
| *D. acanthoptera* | Infrequent[3] | 6[4] | 137.1[5] | 1.21[4] | 7.84[6] | 1.13[4] | 8.20[7] | 5.83[4] |
| *D. pachea* | Rapid[3] | 3[4] | 39.5[5] | 1.12[4] | 19.87[6] | 1.02[4] | 22.75[7] | 16.53[4] |
| *D. nannoptera* | Rapid[3] | 4[4] | 5.1[5] | 1.08[4] | 20.89[6] | 0.99[4] | 19.81[7] | 15.74[4] |
| *D. silvestris* | Infrequent[3] | 21[5] | -- | 3.16[5] | -- | 3.24[2] | -- | -- |
| *D. guttifera* | Rapid[3] | 5[4] | 7.8[5] | 0.99[4] | 17.77[6] | 0.94[4] | -- | 10.29[4] |
| *D. funebris* | Infrequent[5] | -- | 16.9[5] | 1.41[8] | 15.84[9] | 1.36[2] | 10.65[9] | 8.29[9] |
| *D. arawakana* | Infrequent | 5 | 30.4 | 0.889 | 3.291 | 0.798 | 2.408 | 1.471 |
| *D. immigrans* | -- | -- | 51.7 | 1.373 | 3.378 | 1.191 | 3.175 | 1.484 |

Sources: [1] Holman et al. 2008; [2] Interpolated from female thorax length; [3] Markow 2002; [4] Pitnick et al. 1995; [5] Markow 1996; [6] Pitnick et al. 1999; [7] Pitnick 1996; [8] Atkinson 1979; [9] Joly and Bressac 1994.

**Table S5.** Best-fitting models of continuous trait evolution and their corresponding estimates for rates of trait evolution and root states. We compared Brownian motion, early-burst, and Ornstein-Uhlenbeck models. All variables were log-transformed for analysis, but root state estimates have been back-transformed to simplify interpretation.

| Trait | N of species | Model of evolution | Rate estimate ($\sigma^2$) | Root state estimate ($z_0$) |
|---|---|---|---|---|
| **Age of female reproductive maturity (days)** | 29 | Ornstein-Uhlenbeck | 0.120 | 4.918 |
| **Mating latency (min)** | 15 | Ornstein-Uhlenbeck | 0.187 | 12.743 |
| **Mating duration (min)** | 30 | Ornstein-Uhlenbeck | 0.245 | 10.206 |
| **Female remating rate (matings/day)** | 15 | Brownian motion | 0.093 | 0.268 |
| **Female thorax length (mm)** | 31 | Brownian motion | 0.003 | 1.120 |
| **SR length (mm)** | 30 | Brownian motion | 0.069 | 4.121 |
| **ST area ($mm^2$)** | 15 | Ornstein-Uhlenbeck | 0.094 | 0.006 |
| **ST pigment (grayvalue)** | 15 | Ornstein-Uhlenbeck | 0.016 | 96.834 |
| **Non-RT female body mass (mg)** | 15 | Brownian motion | 0.011 | 0.324 |
| **FRT mass (mg)** | 15 | Ornstein-Uhlenbeck | 0.138 | 0.092 |
| **Male thorax length (mm)** | 31 | Brownian motion | 0.002 | 0.929 |
| **Sperm length (mm)** | 30 | Brownian motion | 0.059 | 2.132 |
| **Testis length (mm)** | 21 | Brownian motion | 0.038 | 3.384 |
| **Non-RT male body mass (mg)** | 15 | Brownian motion | 0.012 | 0.218 |
| **MRT mass (mg)** | 15 | Brownian motion | 0.006 | 0.028 |

**Table S6.** Best-fitting M*k* models of discrete trait evolution and their corresponding transition rate estimates. We compared equal-rates, symmetric transition, and all-rates-different models.

| Trait | N of species | Model of evolution | Transition rate estimate ($q$) |
|---|---|---|---|
| **Female remating rate (rapid or infrequent)** | 30 | Equal-rates | 0.024 |
| **Testis color** | 15 | Equal-rates | 0.007 |

**Table S7.** Phylogenetic generalized least squares (PGLS) regressions quantifying relationships among reproductive traits in *Drosophila* species. n = 15 species for all models except those that include sperm length, for which n = 10. All continuous variables were log-transformed. Coefficients are provided for the predictor of interest, when applicable; coefficients for the size covariates included in all models are presented in Table S11.

| Dependent variable | Predictor of interest | Size covariate | Value | Std error | *t* | *p* |
|---|---|---|---|---|---|---|
| **SR length** | Female thorax length | N/A | 2.087 | 1.421 | 1.468 | 0.166 |
| **SR length** | Age of female reproductive maturity | Female thorax length | 0.296 | 0.279 | 1.064 | 0.308 |
| **SR length** | Female remating rate | Female thorax length | 0.018 | 0.166 | 0.106 | 0.917 |
| **SR length** | MRT mass | Female thorax length | 0.262 | 0.662 | 0.396 | 0.699 |
| **SR length** | Sperm length | Female thorax length | 0.895 | 0.143 | 6.280 | <0.001 *** |
| **SR length** | Testis length | Female thorax length | 1.235 | 0.134 | 9.193 | <0.001 *** |
| **ST area** | Female thorax length | N/A | 4.335 | 1.367 | 3.170 | 0.007 ** |
| **ST area** | Age of female reproductive maturity | Female thorax length | 0.239 | 0.283 | 0.846 | 0.414 |
| **ST area** | Female remating rate | Female thorax length | -0.206 | 0.155 | -1.322 | 0.211 |
| **ST area** | MRT mass | Female thorax length | -0.412 | 0.607 | -0.679 | 0.510 |
| **ST pigmentation** | Female non-RT body mass | N/A | -0.095 | 0.195 | -0.486 | 0.635 |
| **ST pigmentation** | Age of female reproductive maturity | Female non-RT body mass | 0.142 | 0.155 | 0.920 | 0.376 |
| **ST pigmentation** | Female remating rate | Female non-RT body mass | -0.096 | 0.077 | -1.246 | 0.237 |
| **ST pigmentation** | MRT mass | Female non-RT body mass | -0.287 | 0.290 | -0.988 | 0.343 |
| **FRT mass** | Female non-RT body mass | N/A | 0.216 | 0.026 | 8.472 | <0.001 *** |
| **FRT mass** | Age of female reproductive maturity | Female non-RT body mass | 0.078 | 0.208 | 0.374 | 0.715 |
| **FRT mass** | Female remating rate | Female non-RT body mass | 0.092 | 0.086 | 1.067 | 0.307 |
| **FRT mass** | MRT mass | Female non-RT body mass | 0.372 | 0.261 | 1.426 | 0.179 |

| | | | | | | |
|---|---|---|---|---|---|---|
| **Female remating rate** | Female non-RT body mass | N/A | 0.497 | 0.706 | 0.704 | 0.494 |
| **Female remating rate** | Age of female reproductive maturity | Female non-RT body mass | -0.942 | 0.442 | -2.132 | 0.054 |
| **Female remating rate** | MRT mass | Female non-RT body mass | 1.864 | 0.771 | 2.418 | 0.033 * |
| **Testis length** | Male thorax length | N/A | 3.153 | 0.923 | 3.417 | 0.005 ** |
| **Testis length** | Female remating rate | Male thorax length | -0.038 | 0.114 | -0.333 | 0.745 |
| **Testis length** | MRT mass | Male thorax length | 0.074 | 0.459 | 0.161 | 0.875 |
| **Sperm length** | Male thorax length | N/A | 2.356 | 1.250 | 1.885 | 0.096 |
| **Sperm length** | Female remating rate | Male thorax length | -0.052 | 0.135 | -0.389 | 0.709 |
| **Sperm length** | MRT mass | Male thorax length | -0.271 | 0.551 | -0.492 | 0.638 |
| **MRT mass** | Male non-RT body mass | N/A | 0.212 | 0.194 | 1.096 | 0.293 |
| **MRT mass** | Female remating rate | Male non-RT body mass | 0.099 | 0.079 | 1.254 | 0.234 |
| **Mating latency** | Female non-RT body mass | N/A | 1.083 | 0.052 | 20.795 | <0.001 *** |
| **Mating latency** | Age of female reproductive maturity | Female non-RT body mass | 0.435 | 0.294 | 1.482 | 0.164 |
| **Mating latency** | Female remating rate | Female non-RT body mass | -0.024 | 0.137 | -0.173 | 0.866 |
| **Mating latency** | MRT mass | Female non-RT body mass | -0.338 | 0.411 | -0.821 | 0.428 |
| **Mating duration** | Female non-RT body mass | N/A | -0.693 | 0.375 | -1.847 | 0.088 |
| **Mating duration** | Age of female reproductive maturity | Female non-RT body mass | 0.233 | 0.237 | 0.983 | 0.345 |
| **Mating duration** | Female remating rate | Female non-RT body mass | -0.052 | 0.141 | -0.370 | 0.718 |
| **Mating duration** | MRT mass | Female non-RT body mass | -0.661 | 0.496 | -1.334 | 0.207 |

**Table S8.** Phylogenetic generalized least squares (PGLS) regressions quantifying relationships among reproductive traits in expanded dataset. All continuous variables were log-transformed. Coefficients are provided for the predictor of interest, when applicable; coefficients for the size covariates included in all models are presented in Table S12.

| Dependent variable | N of species | Predictor of interest | Size covariate | Value | Std error | *t* | *p* |
|---|---|---|---|---|---|---|---|
| **SR length** | 30 | Female thorax length | N/A | 1.419 | 0.871 | 1.630 | 0.115 |
| **SR length** | 29 | Age of female reproductive maturity | Female thorax length | 0.224 | 0.319 | 0.765 | 0.451 |
| **SR length** | 30 | Sperm length | Female thorax length | 0.983 | 0.077 | 12.791 | <0.001 *** |
| **SR length** | 20 | Testis length | Female thorax length | 1.234 | 0.152 | 8.122 | <0.001 *** |

**Table S9.** Coefficients from a phylogenetic ANCOVA that tests for the effect of testis color on female remating rate, including male body mass as a size covariate. All continuous variables were log-transformed.

| Dependent variable | Model component | DF | *F* | *p* |
|---|---|---|---|---|
| Female remating rate | Intercept | 1 | 2.307 | 0.157 |
| | Male body mass | 1 | 3.361 | 0.094 |
| | Testis color | 2 | 0.789 | 0.478 |

**Table S10.** Multivariate Brownian evolution models (Trait 1 = sperm length, Trait 2 = SR length, n = 29 species) with multiple evolutionary correlations. We sought to test whether rates of SR or sperm length evolution (or their coevolution) differed on branches of the phylogeny where female remating is reconstructed as either rapid or infrequent (as determined by stochastic mapping of ancestral states for this discrete trait; see Figure 4). Low AIC scores indicate a better model fit; scores within two units of each other indicate similar support for the models under comparison (Burham and Anderson 1994).

| **Model 1: common rates, common correlation** | | | | |
|---|---|---|---|---|
| | R[1,1] | R[1,2] | R[2,2] | AIC |
| **Fitted** | 0.059 | 0.060 | 0.071 | 97.333 |
| **Model 2: different rates, common correlation** | | | | |
| | R[1,1] | R[1,2] | R[2,2] | AIC |
| **Infrequent** | 0.048 | 0.044 | 0.047 | 98.006 |
| **Rapid** | 0.072 | 0.077 | 0.096 | |

| | | | | |
|---|---|---|---|---|
| **Model 2b: different rates (sperm length only), common correlation** | | | | |
| | R[1,1] | R[1,2] | R[2,2] | AIC |
| **Infrequent** | 0.064 | 0.062 | 0.071 | 99.112 |
| **Rapid** | 0.056 | 0.058 | 0.071 | |
| **Model 2c: different rates (SR length only), common correlation** | | | | |
| | R[1,1] | R[1,2] | R[2,2] | AIC |
| **Infrequent** | 0.059 | 0.052 | 0.055 | 96.965 |
| **Rapid** | 0.059 | 0.065 | 0.084 | |
| **Model 3: common rates, different correlation** | | | | |
| | R[1,1] | R[1,2] | R[2,2] | AIC |
| **Infrequent** | 0.062 | 0.067 | 0.069 | 97.345 |
| **Rapid** | 0.062 | 0.058 | 0.069 | |
| **Model 3b: different rates (sperm length only), different correlation** | | | | |
| | R[1,1] | R[1,2] | R[2,2] | AIC |
| **Infrequent** | 0.070 | 0.067 | 0.071 | 98.643 |
| **Rapid** | 0.055 | 0.055 | 0.071 | |
| **Model 3c: different rates (SR length only), different correlation** | | | | |
| | R[1,1] | R[1,2] | R[2,2] | AIC |
| **Infrequent** | 0.059 | 0.055 | 0.057 | 97.728 |
| **Rapid** | 0.059 | 0.062 | 0.081 | |
| **Model 4: no common structure** | | | | |
| | R[1,1] | R[1,2] | R[2,2] | AIC |
| **Infrequent** | 0.053 | 0.049 | 0.052 | 99.550 |
| **Rapid** | 0.065 | 0.068 | 0.087 | |

**Table S11.** Coefficients for each size covariate from the corresponding phylogenetic generalized least squares (PGLS) regressions shown in Table S7. All continuous variables were log-transformed. λ values represent the degree of phylogenetic dependence for each variable, as estimated using maximum likelihood.

| **Dependent variable** | **Size covariate** | **Predictor of interest** | **Value** | **Std error** | ***t*** | ***p*** | **λ** |
|---|---|---|---|---|---|---|---|
| **SR length** | Female thorax length | -- | 2.087 | 1.421 | 1.468 | 0.166 | 0.999 |
| **SR length** | Female thorax length | Age of female reproductive maturity | 2.513 | 1.465 | 1.715 | 0.112 | 0.998 |
| **SR length** | Female thorax length | Female remating rate | 2.075 | 1.520 | 1.366 | 0.197 | 0.998 |

| | | | | | | | |
|---|---|---|---|---|---|---|---|
| **SR length** | Female thorax length | MRT mass | 1.935 | 1.555 | 1.244 | 0.237 | 0.996 |
| **SR length** | Female thorax length | Sperm length | -1.118 | 0.717 | -1.560 | 0.163 | 1.038 |
| **SR length** | Female thorax length | Testis length | -2.109 | 0.720 | -2.929 | 0.013 * | 0.955 |
| **ST area** | Female thorax length | -- | 4.335 | 1.367 | 3.170 | 0.007 ** | 0.973 |
| **ST area** | Female thorax length | Age of female reproductive maturity | 4.642 | 1.425 | 3.257 | 0.007 ** | 0.972 |
| **ST area** | Female thorax length | Female remating rate | 4.890 | 1.356 | 3.607 | 0.004 ** | 0.954 |
| **ST area** | Female thorax length | MRT mass | 4.922 | 1.465 | 3.360 | 0.006 ** | 0.928 |
| **ST pigmentation** | Female non-RT body mass | -- | -0.095 | 0.195 | -0.486 | 0.635 | 0.841 |
| **ST pigmentation** | Female non-RT body mass | Age of female reproductive maturity | -0.081 | 0.195 | -0.414 | 0.686 | 0.780 |
| **ST pigmentation** | Female non-RT body mass | Female remating rate | -0.027 | 0.201 | -0.133 | 0.897 | 0.909 |
| **ST pigmentation** | Female non-RT body mass | MRT mass | 0.000 | 0.215 | 0.000 | 0.999 | 0.936 |
| **FRT mass** | Female non-RT body mass | -- | 0.216 | 0.026 | 8.472 | <0.001 *** | -0.255 |
| **FRT mass** | Female non-RT body mass | Age of female reproductive maturity | 0.122 | 0.220 | 1.037 | 0.320 | 0.122 |
| **FRT mass** | Female non-RT body mass | Female remating rate | 0.146 | 0.195 | 0.746 | 0.470 | -0.032 |
| **FRT mass** | Female non-RT body mass | MRT mass | 0.008 | 0.225 | 0.036 | 0.972 | -0.087 |
| **Female remating rate** | Female non-RT body mass | -- | 0.497 | 0.706 | 0.704 | 0.494 | 0.897 |
| **Female remating rate** | Female non-RT body mass | Age of female reproductive maturity | 0.193 | 0.661 | 0.292 | 0.775 | 0.949 |
| **Female remating rate** | Female non-RT body mass | MRT mass | -0.093 | 0.648 | -0.144 | 0.888 | 0.635 |
| **Testis length** | Male thorax length | -- | 3.153 | 0.923 | 3.417 | 0.005 ** | 1.013 |
| **Testis length** | Male thorax length | Female remating rate | 3.253 | 0.968 | 3.360 | 0.006 ** | 1.010 |
| **Testis length** | Male thorax length | MRT mass | 3.178 | 0.980 | 3.242 | 0.007 ** | 1.009 |
| **Sperm length** | Male thorax length | -- | 2.356 | 1.250 | 1.885 | 0.096 | 1.045 |
| **Sperm length** | Male thorax length | Female remating rate | 0.989 | 1.663 | 0.595 | 0.571 | 1.007 |

| | | | | | | | |
|---|---|---|---|---|---|---|---|
| **Sperm length** | Male thorax length | MRT mass | 0.832 | 1.664 | 0.500 | 0.633 | 1.009 |
| **MRT mass** | Male non-RT body mass | -- | 0.212 | 0.194 | 1.096 | 0.293 | 0.982 |
| **MRT mass** | Male non-RT body mass | Female remating rate | 0.121 | 0.219 | 0.551 | 0.592 | 0.960 |
| **Mating latency** | Female non-RT body mass | -- | 1.083 | 0.052 | 20.795 | <0.001 *** | -0.831 |
| **Mating latency** | Female non-RT body mass | Age of female reproductive maturity | 0.676 | 0.308 | 2.197 | 0.048 * | -0.237 |
| **Mating latency** | Female non-RT body mass | Female remating rate | 1.138 | 0.293 | 3.888 | 0.002 ** | -0.783 |
| **Mating latency** | Female non-RT body mass | MRT mass | 1.632 | 0.293 | 5.576 | <0.001 *** | -1.320 |
| **Mating duration** | Female non-RT body mass | -- | -0.693 | 0.375 | -1.847 | 0.088 | 0.993 |
| **Mating duration** | Female non-RT body mass | Age of female reproductive maturity | -0.607 | 0.386 | -1.574 | 0.142 | 0.993 |
| **Mating duration** | Female non-RT body mass | Female remating rate | -0.683 | 0.390 | -1.754 | 0.105 | 0.977 |
| **Mating duration** | Female non-RT body mass | MRT mass | -0.611 | 0.373 | -1.637 | 0.128 | 0.899 |

**Table S12.** Coefficients for each size covariate from the corresponding phylogenetic generalized least squares (PGLS) regressions shown in Table S8. All continuous variables were log-transformed. λ values represent the degree of phylogenetic dependence for each variable, as estimated using maximum likelihood.

| **Dependent variable** | **N of species** | **Size covariate** | **Predictor of interest** | **Value** | **Std error** | ***t*** | ***p*** | **λ** |
|---|---|---|---|---|---|---|---|---|
| **SR length** | 30 | Female thorax length | -- | 1.419 | 0.871 | 1.630 | 0.115 | 0.980 |
| **SR length** | 29 | Female thorax length | Age of female reproductive maturity | -0.493 | 1.427 | -0.345 | 0.733 | 0.944 |
| **SR length** | 30 | Female thorax length | Sperm length | -0.685 | 0.470 | -1.458 | 0.157 | 0.849 |
| **SR length** | 20 | Female thorax length | Testis length | -1.556 | 0.803 | -1.939 | 0.069 | 0.803 |

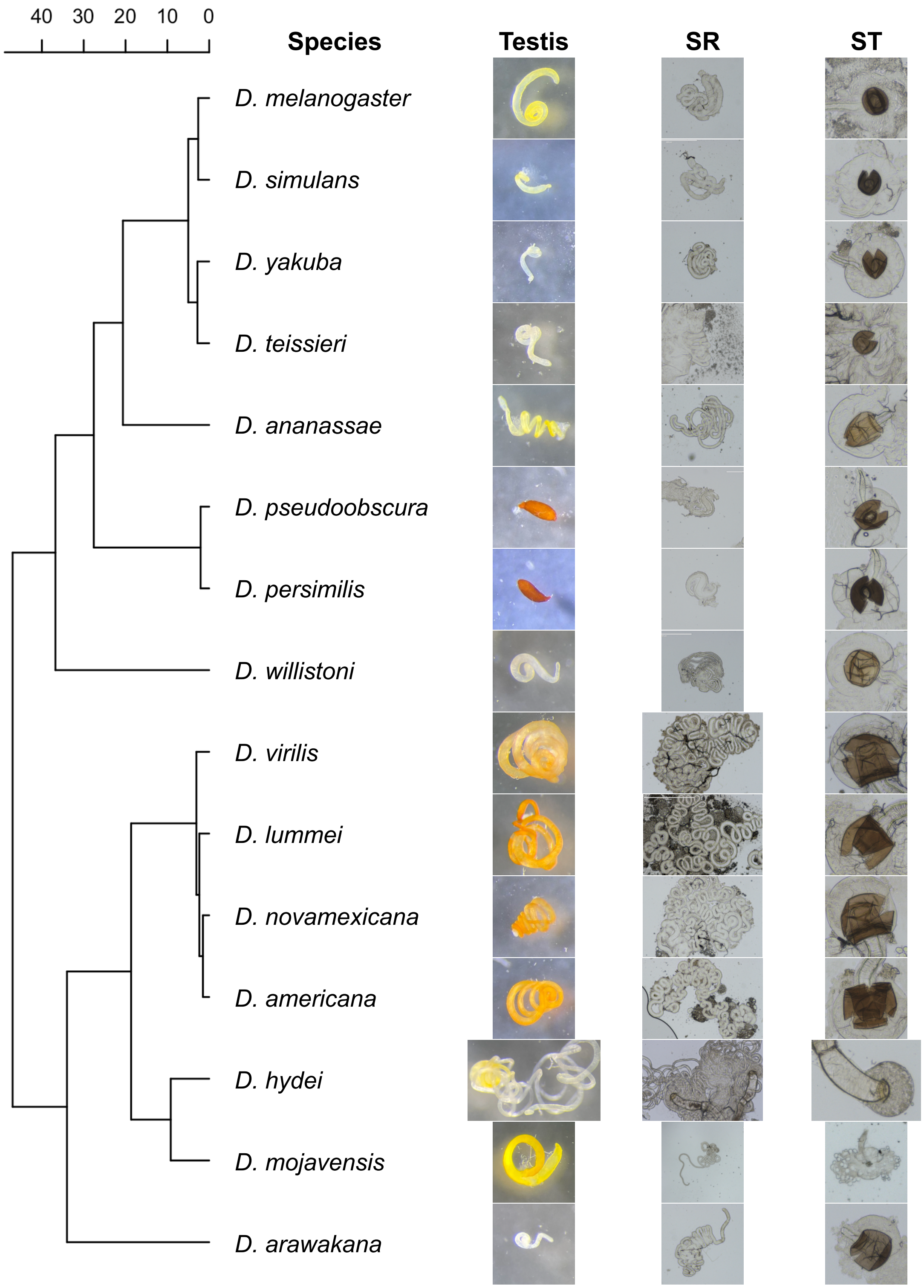

**Figure S1.** Diversity in reproductive morphology across 15 *Drosophila* species. Columns, from left to right, show photos of testes, seminal receptacles (SR), and spermathecae (ST) from each species. Scales are consistent within a trait, but not between traits.

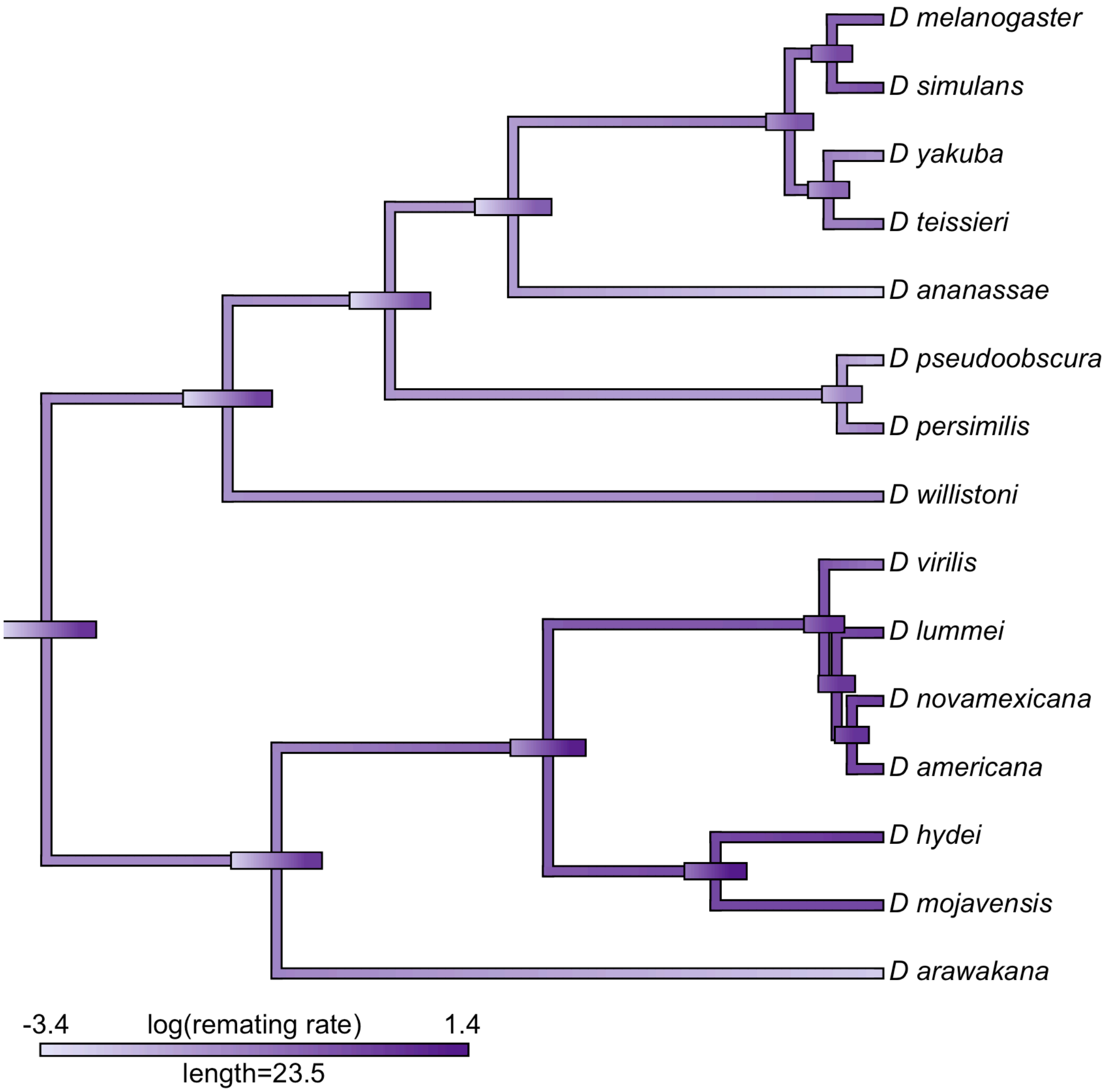

**Figure S2.** A projection of our directly observed, quantitative estimate of remating rate (on a log-scale) onto the *Drosophila* phylogeny. The color gradient shows observed (at the tips) or estimated trait values. Error bars show the uncertainty associated with the estimated values for the trait at internal nodes of the tree.

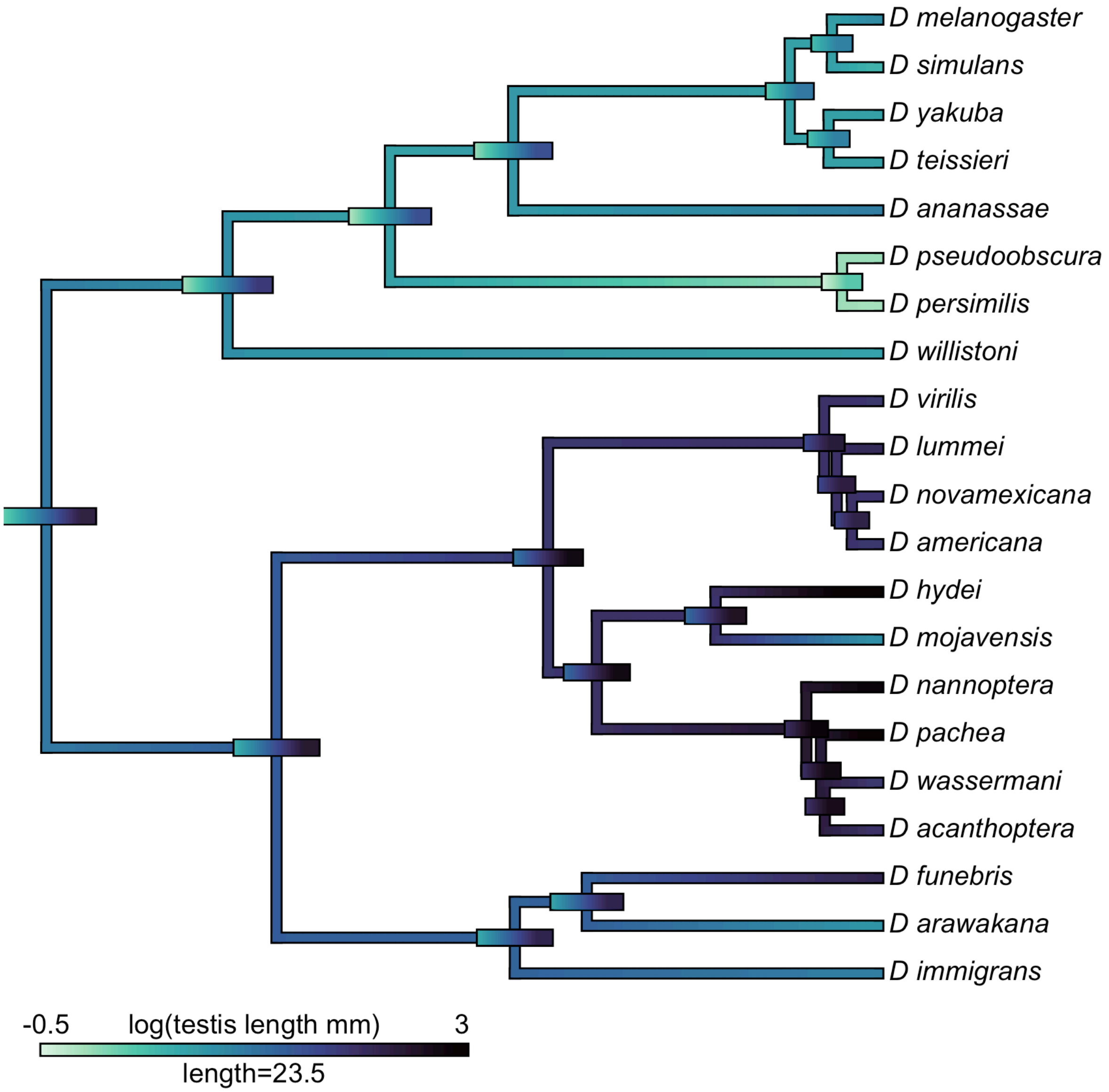

**Figure S3.** A projection of testis length (on a log-scale) onto the *Drosophila* phylogeny. The color gradient shows observed (at the tips) or estimated trait values. Error bars show the uncertainty associated with the estimated values for the trait at internal nodes of the tree.

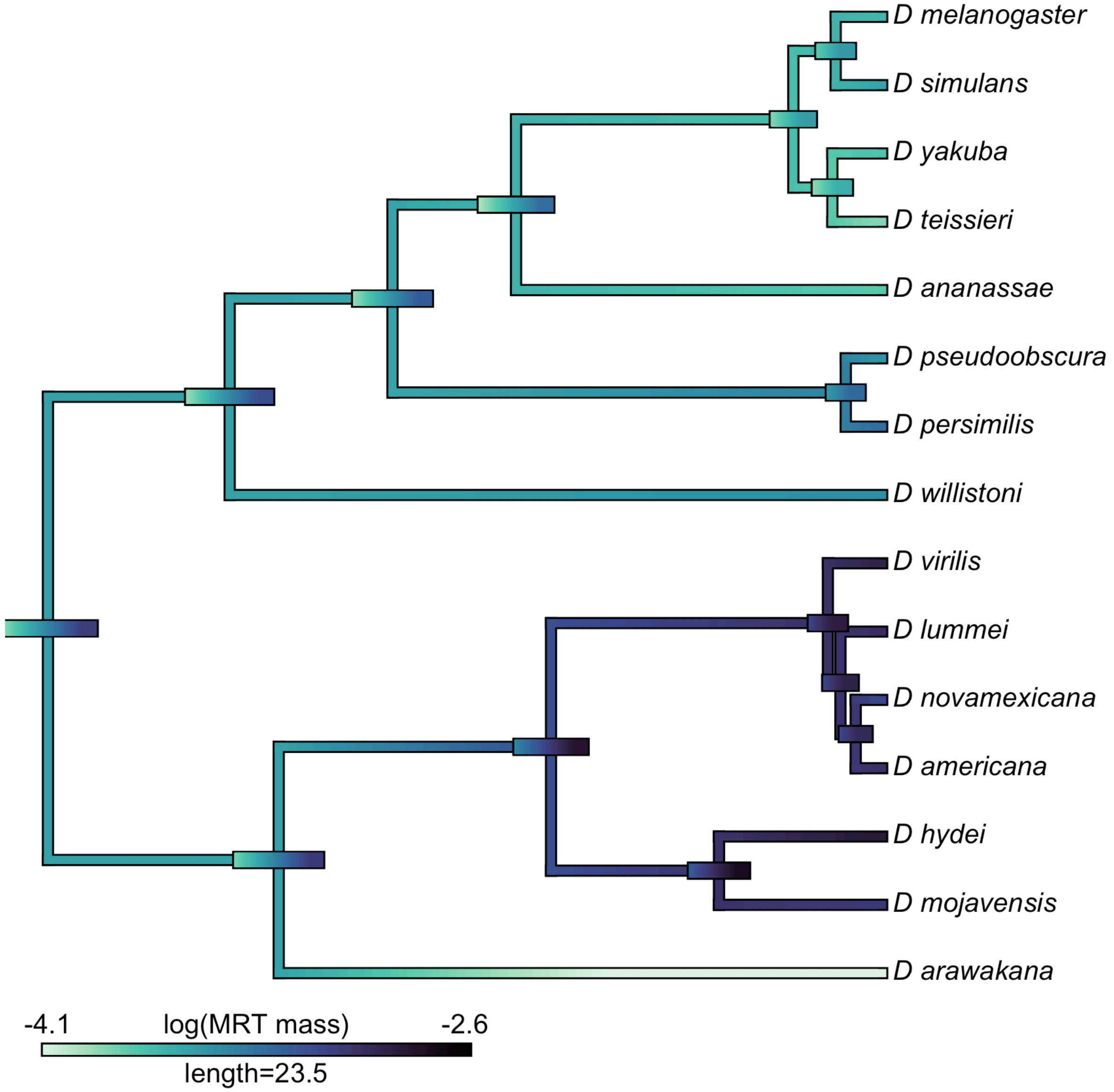

**Figure S4.** A projection of MRT mass (on a log-scale) onto the *Drosophila* phylogeny. The color gradient shows observed (at the tips) or estimated trait values. Error bars show the uncertainty associated with the estimated values for the trait at internal nodes of the tree.

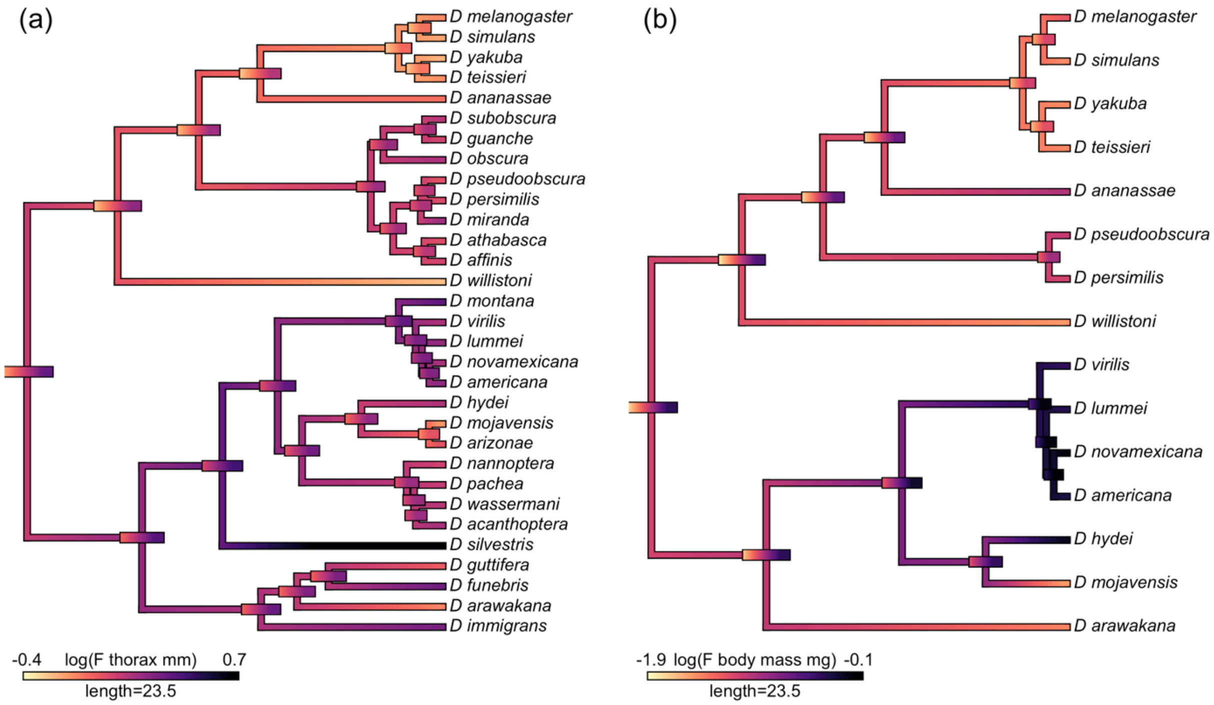

**Figure S5.** A projection of (a) female thorax length and (b) non-RT female body mass (both on a log-scale) onto the *Drosophila* phylogeny. The color gradient shows observed (at the tips) or estimated (internal branches) trait values. Error bars show the uncertainty associated with the estimated values for the trait at internal nodes of the tree.

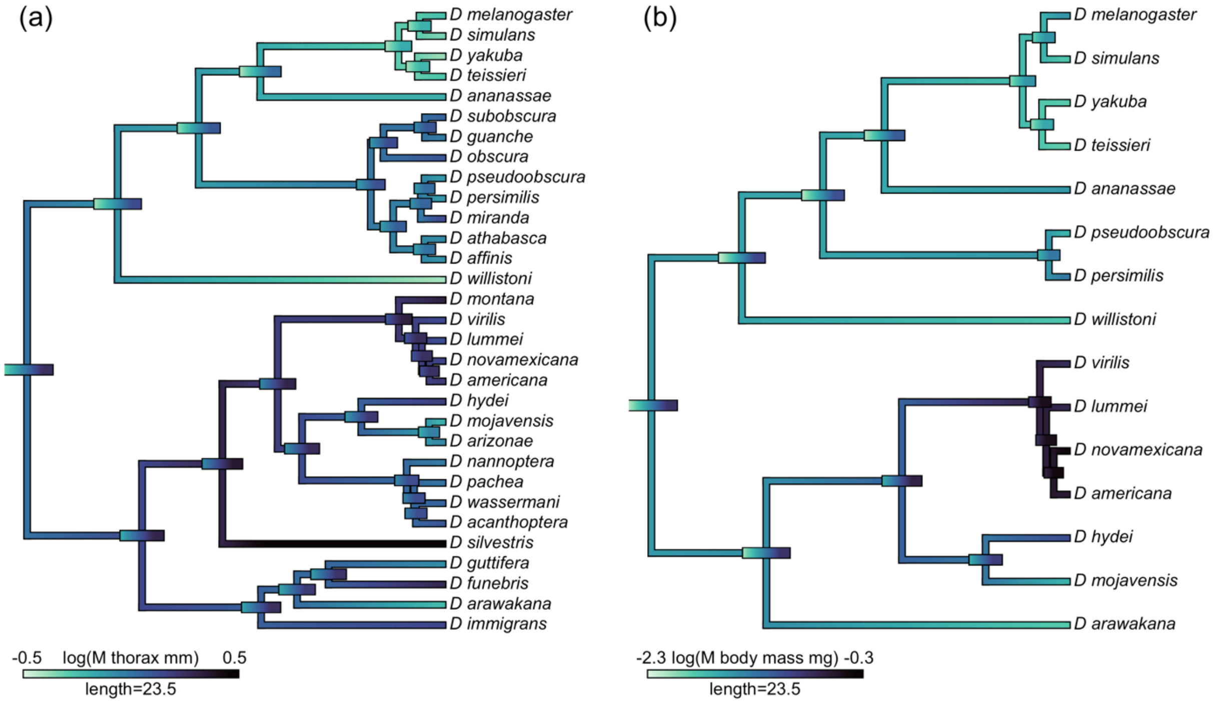

**Figure S6.** A projection of (a) male thorax length and (b) non-RT male body mass (both on a log-scale) onto the *Drosophila* phylogeny. The color gradient shows observed (at the tips) or estimated trait values. Error bars show the uncertainty associated with the estimated values for the trait at internal nodes of the tree.

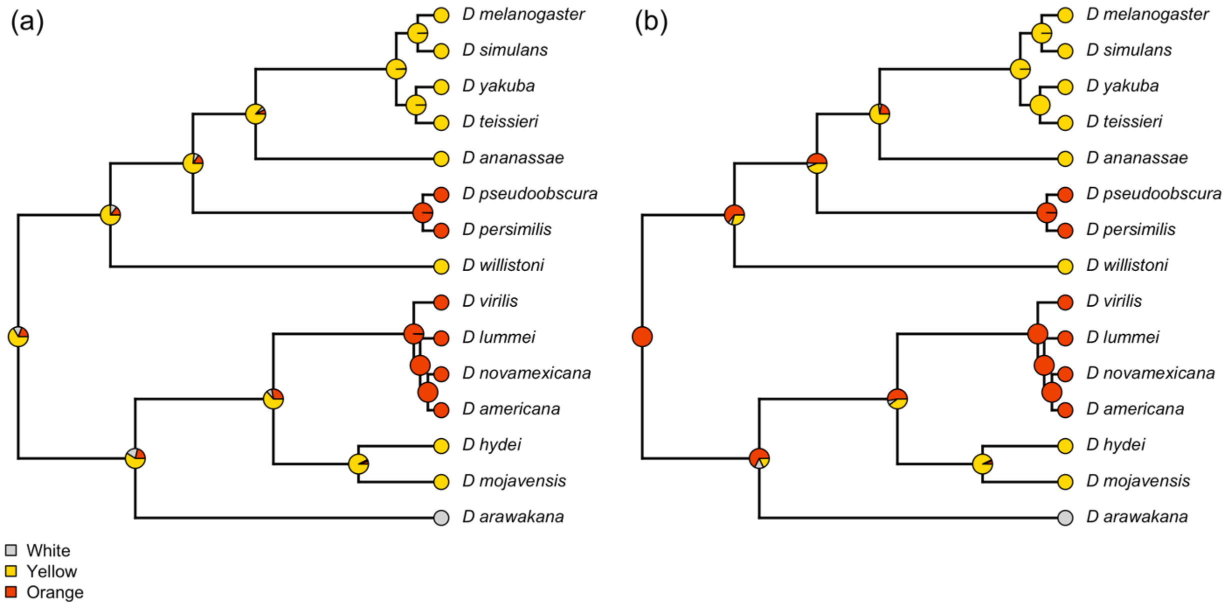

**Figure S7.** Posterior probabilities for testis color at nodes from 1000 stochastic character maps. (a) depicts the posterior probabilities as estimated with no root node state specified, while (b) depicts the posterior probabilities when the root node is constrained to orange.

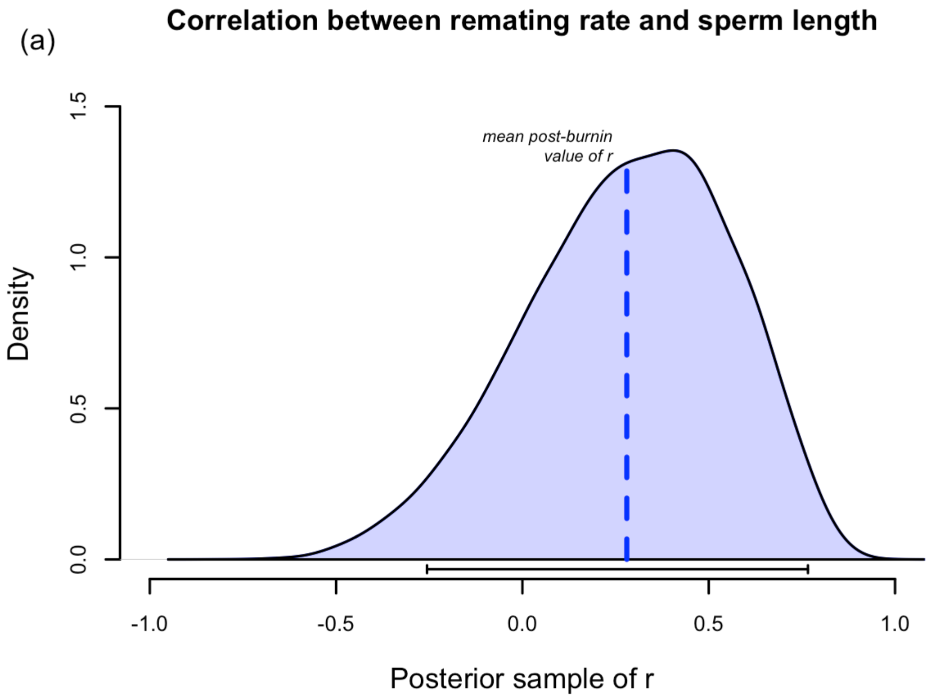

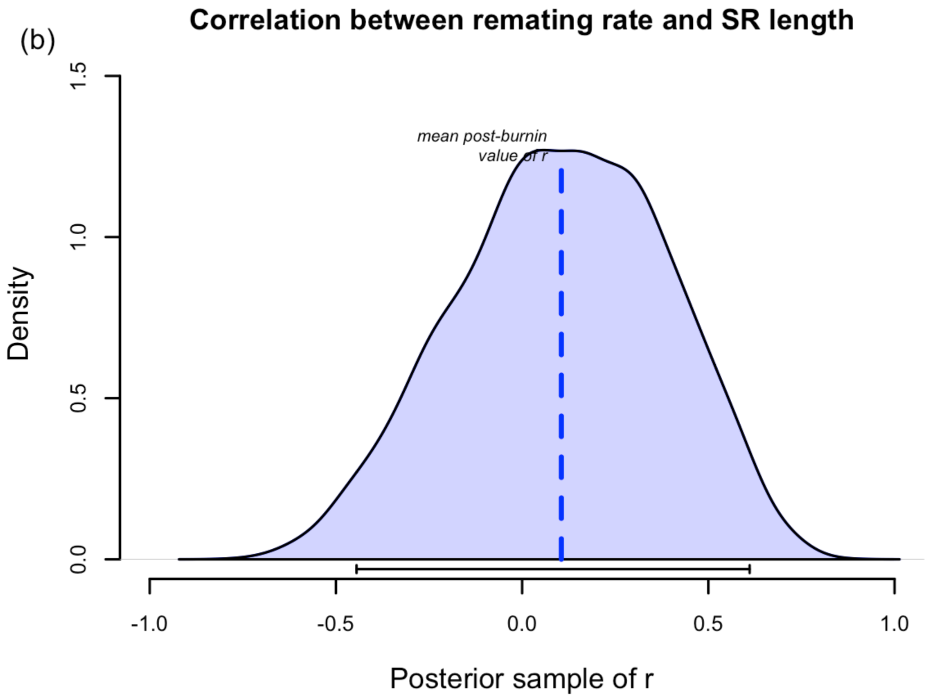

**Figure S8.** Posterior density of the correlation between female remating rate (rapid or infrequent) and (a) sperm length or (b) SR length. Data from 29 species are used in both analyses. Bars along the x-axes indicate 95% high-probability density (HPD) intervals. We found no significant correlation between female remating rate and either sperm or SR length, as indicated by the 95% high-probability density (HPD) interval overlapping with zero.

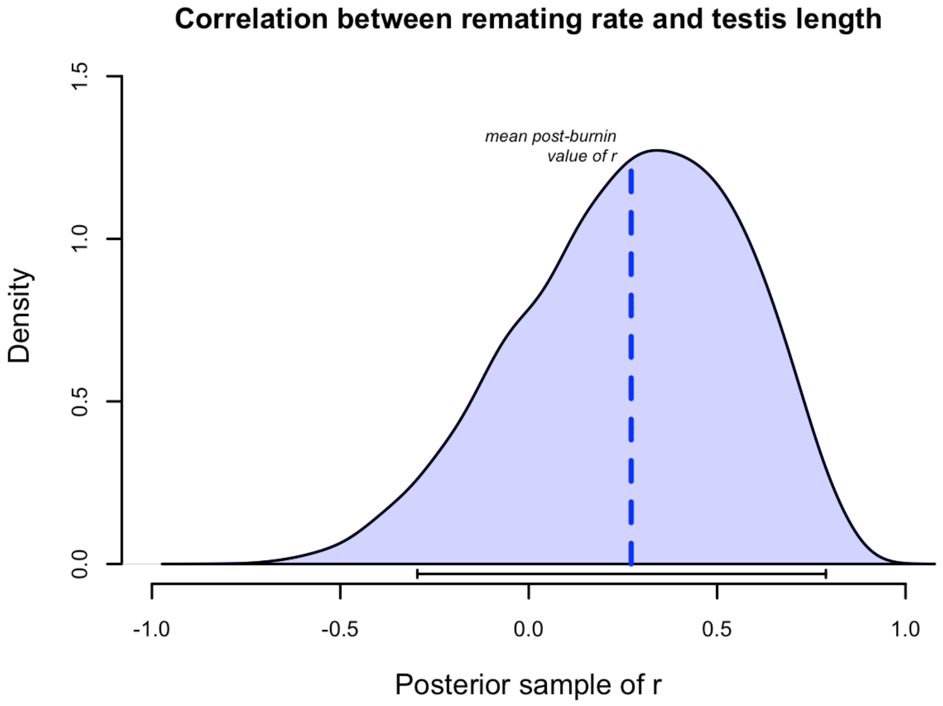

**Figure S9.** Posterior density of the correlation between female remating rate (rapid or infrequent) and testis length. Data from 20 species are used in this analysis. The bar along the x-axis indicates the 95% high-probability density (HPD) interval. We found no significant correlation between female remating rate and testis length, as indicated by the 95% high-probability density (HPD) interval overlapping with zero.

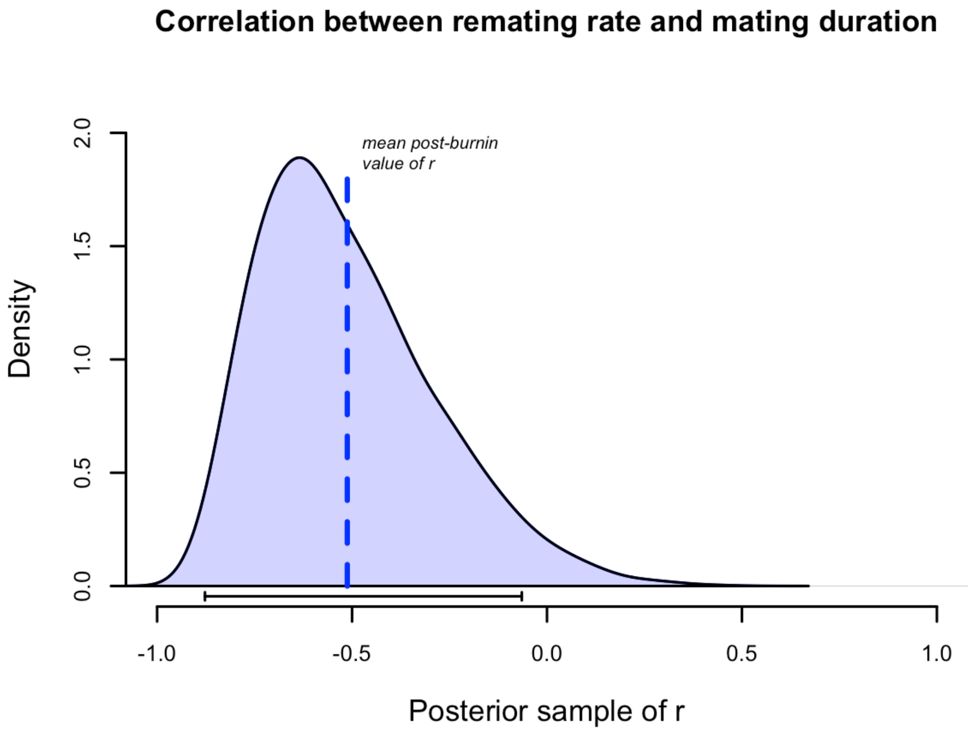

**Figure S10.** Posterior density of the correlation between female remating rate (rapid or infrequent) and mating duration. Data from 29 species were used in this analysis. The bar along the x-axis indicates the 95% high-probability density (HPD) interval. We found a significant negative correlation between female remating rate and mating duration as indicated by the 95% high-probability density (HPD) interval not overlapping with zero.
